# High-resolution and quantitative imaging of the post mortem brain

**DOI:** 10.64898/2026.01.18.700174

**Authors:** A.-M. Oros-Peusquens, N.J. Shah

**Affiliations:** Institute of Neuroscience and Medicine - 4, Research Centre Juelich, 52425 Juelich, Germany; Department of Neurology, Faculty of Medicine, JARA, RWTH Aachen University, 52074 Aachen, Germany

## Abstract

MRI of fixed tissue is an excellent way to study pathological changes caused by different diseases with great anatomical detail. It is, however, known that properties of tissue change with fixation. The aim of this study was to determine the variability of several quantitative MRI (qMRI) parameters in fixed brain tissue obtained from donors unaffected by neurological conditions and investigate the existence of quantitative parameters which vary little between specimens. We introduce a 3D method for high-resolution mapping of water content, T_1_ and T_2_* relaxation times and parameters characterising magnetisation transfer and apply it at 3T to 7 whole, fixed human brains (3 male, 4 female, aged between 47 and 79 years, mean age 67 years). The qMRI parameters determined include relaxation rates T_1_ and T_2_*, MT ratio and T_1_ and T_2_* after MT. From these we can further derive semiquantitative MT parameters such as the exchange rate (k_trans_) and bound pool fraction (f_bound_). Correlations between these parameters are investigated. In addition, truly quantitative water content determined non-invasively with MRI is reported on whole human post mortem brains – to our knowledge, for the first time. Water content was found to have mean values of 73% for WM and 85% for GM with standard deviation below 2.5% over 7 brains, and thus a few percent units higher than *in vivo* (69% and 81%) and of comparable constancy.

## Introduction

The past decade has brought very high field imagining nearly in the realm of routine. Perhaps through the realisation that MRI at very high fields can become a method for microscopy and even an alternative to histology, increasing interest and effort has been dedicated to post mortem brain imaging. However, while the anatomy and connectivity of well fixed specimens still correspond to those of the living brains, the MR properties are changed by fixation in a not fully understood way. This is perhaps not surprising given that the distribution of intra/extracellular space and material in tissue 4is modified by fixation [Fox 1985, Torack 1965, Cragg 1979].

First information about relaxation properties of fixed brains emerged early on (Thickman 1983, Tovi and Ericsson 1992, Blamire et al. 1999) but fixed brain imaging remained for quite some time a sparsely populated niche, mainly used to exemplify the power of medium and high field imaging systems (Pfefferbaum 2004, Augustinack 2005).

Fixed tissue has altered T_1_ and T_2_ compared to in-vivo. Values for T_2_ relaxation of formaldehyde-fixed brain tissue were reported to be T_2_ = 35–70 ms at 0.5–4.7 T (Tovi and Ericsson, 1992, Pfefferbaum et al., 2004, Yong-Hing et al., 2005, D’Arceuil et al., 2007, McNab et al., 2009) and are shortened by roughly a factor of 2 with respect to fresh tissue. Rehydrating fixed tissue in phosphate buffer solution significantly increases T_2_ (Thelwall et al., 2006, Purea et al. 2006, D’Arceuil et al., 2007, Shepherd et al., 2009), potentially even exceeding in-vivo values (Shepherd et al., 2009). These results suggest that short T_2_ is driven, at least in part, by the presence of fixative in tissue. A practical consequence of the shortened T_2_ is that it makes diffusion studies with manufacturer-provided diffusion sequences (typically double-refocused spin echo with EPI read-out and TE>100ms) unfeasible. Alternative methods have mostly involved the use of animal scanners with high gradient strength and/or slower diffusion acquisition methods [d’Arceuil et al., 2007, McNab et al., 2009, Miller et al. 2011]. Several studies have found reduced T_1_ values of 300–500 ms at field strengths of 0.5–4.7 T (Tovi and Ericsson 1992, Pfefferbaum et al., 2004 and Yong-Hing et al., 2005, d’Arceuil et al., 2007, McNab et al., 2009). Rehydration in buffer solution does not affect T_1_ much ( D’Arceuil et al., 2007 and Shepherd et al., 2009). For imaging purposes, this is in general an advantage, allowing for shorter repetition times than *in vivo* or higher SNR.

At very high fields, fiber orientation adds another dimension to the traditional MRI contrast (Duyn et al., 2007, Lee et al., 2011, Wharton and Bowtell, 2012). These effects can be elegantly studied in fixed brains, which can be, for example, easily reoriented with respect to the magnetic field. However, at 3T these effects are still reduced.

Despite the fact that all MR imaging included the proton density as a fundamental parameter, to our knowledge, reports of truly quantitative water content measured by MRI did not exist for whole fixed brains before the present study.

The aim of this study was to investigate post mortem formalin-fixed brains quantitatively in order to provide a baseline for MR parameters characteristic of healthy brain tissue, in particular regarding their water content. A 3D quantitative imaging protocol was developed and optimised for the range of parameters met in fixed tissue. It is a 2-point mapping method based on a similar principle to the ones presented by Deoni [Deoni et al. 2005] and Helms et al. [Helms et al. 2008] but differing substantially from previous work in the details and optimisation. The parameters of the 2-point acquisition were optimised with respect to the flip angle, to ensure good accuracy and precision of the two-point mapping method. Relaxation times (T_1_, T_2_*) and equilibrium magnetisation density (M_z_ ^0^) were determined for 7 whole human brains. The same quantities were measured following a magnetisation transfer (MT) preparation module. From the changes induced by MT, magnetisation transfer ratio (MTR), magnetisation transfer rate (k_trans_) and bound fraction *f_bound_*were determined. Correlations between different parameters were investigated and compared to the same quantities measured *in vivo* with the same method for one volunteer. In addition to providing „hard numbers‟ for the characterisation of fixed tissue, the availability of several quantitative parameters allows for good segmentation of tissue – exemplified on the thalamus.

## Materials and methods

### 2.1 Brains and volunteer

The brains were obtained in accordance with the requirements of the local ethics committees from the brain donor programmes of the Universities of Düsseldorf and Aachen. All donors were free from neurological disorders.

Table 1 summarises information relevant to the properties of tissue, as far as available: gender, age at death, time post mortem, fixation time, fresh weight, weight before MRI.

**Table 1.** a) General information about the brains included in the study. a) General information about MRI examination and observations.

| Brain nr. | Sex | Age | Postmortem time | MRI | Fixation time | Weight [g] |  |
| --- | --- | --- | --- | --- | --- | --- | --- |
|  |  |  |  |  |  | fresh | Formalin |
| 1. | M | 59 | 24 h | 16.07.2010 | 3 ½ mo | 1348 | 1353 |
| 2. | F | 75 | 21 h | 02.07.2010 | 4 mo | 1306 | 1451 |
| 3. | M | 71 | 21 ½ h | 03.09.2010 | 2½ mo | - | 1014 |
| 4. | M | 66 | ~ 24h | 26.01.2010 | 7½ mo | 1279 | 1176 |
| 5. | F | 74 | ~ 24h | 13.01.2010 | 7¾ mo | - | - |
| 6. | F | 79 | 26h | 03.03.2012 | 8 mo | 1197 | 1100 |
| 7. | F | 47 | 15.5h | 03.12.2011 | 13 mo | 1416 | 1300 |
| in vivo | M | 25 | - | 21.05.2010 | - | - | - |

| Brain nr. | Matrix size | Resolution | tissue properties |
| --- | --- | --- | --- |
| 1. | 416 x 512 x 224 | 0.39 x 0.39 x 0.5 |  |
| 2. | 378 x 512 x 284 | 0.39 x 0.39 x 0.6 |  |
| 3. | 416 x 512 x 224 | 0.39 x 0.39 x 0.5 |  |
| 4. | 448 x 512 x 192 | 0.39 x 0.39 x 0.6 | fixation artifact band (T<sub>1</sub>, T<sub>2</sub>*) |
| 5. | 416 x 512 x 192 | 0.39 x 0.39 x 0.6 | fixation artifact: ring (T<sub>1</sub>, T<sub>2</sub>*) |
| 6. | 312 x 384 x 208 | 0.52 x 0.52 x 0.6 |  |
| 7. | 312 x 384 x 208 | 0.52 x 0.52 x 0.6 |  |
| in vivo | 156 x 192 x 128 | 1.04 x 1.04 x 1.0 |  |

The brains were fixed by immersion in 10% neutrally buffered formalin solution (4% formaldehyde concentration). The fixation time amounted to at least 2 ½ months.

For the *in vivo* scan using the same quantitative method, informed consent was obtained from a 26-year old male volunteer, in accordance with the requirements of the local ethics committee.

### 2.2 MRI set-up

A cylindrical plexiglass container was custom made to contain the brains during MRI scanning. Movement was prevented by the inclusion of a variable number of plexiglass plates in the container. The brains were kept in formalin during scanning. A reference probe for water content measurements was also included. It consisted of a 50ml plastic sample tube containing a mixture of 80% H_2_O and 20% D_2_O doped with 70μl Gd-DOTA. The composition was chosen to ensure similar signal intensity (M_0_, T_1_) from the probe to that from the brain.

The open plexiglas container including brain, probe and formalin was kept in a chamber connected to a vacuum pump for 24 hrs prior to scanning in order to eliminate air bubbles.

All measurements were performed using a whole-body 3T scanner (Siemens Tim-Trio), equipped with a gradient coil with maximum field strengths of 40mT/m on each axis. An RF body coil with a homogeneous B_1_-field distribution over the head was used for RF transmit and a 12-element, phased-array head coil was used for signal detection.

As far as possible, the brains were oriented with the cerebellum pointing upwards and the interhemispheric fissure along the long axis of the plexiglas cylinder and parallel to the magnetic field.

### 2.2 MRI protocol

The scanning protocol for quantitative imaging consisted of 2 sets of 2-point 3D mapping protocols, one with and one without magnetisation transfer preparation. The parameters of the acquisitions were optimised separately for post mortem scanning and for *in vivo* parameter mapping, as described in section 2.3.2.

The 3D multiple-echo gradient-echo (3D GRE) sequence offered by the manufacturer was employed. All 3D GRE acquisitions involved in the 2-point method used the same TR of 52 ms and two different flip angles α_1_ = 17° and α_2_ = 75°. Other parameters included: BW = 140Hz/px, TE_1_ = 4.31ms, ΔTE = 8.38ms, RF spoiling, assymetric echo read-out, no parallel imaging. During the standard

GRE acquisitions (without MT preparation) 6 equidistant echoes were read out. In the MT-prepared scans only 4 echoes (same TE_1_ and ΔTE) could be read out for the same TR. The standard MT preparation module was used. Its parameters were: flip angle 500°, pulse duration 10ms, off resonance frequency 1.2 kHz. For each GRE acquisition 2 averages were acquired and the acquisition was repeated 4 times for each set of parameters. Magnitude as well as phase data were saved for each scan.

In addition, one actual flip-angle imaging (AFI) [Yarnykh 2007] acquisition provided B_1_+ maps. AFI is based on the interleaved acquisition of two spoiled FLASH images with the same flip angle and different TRs (TR_1_ and TR_2_). In our case, α=60°, TR_1_/TR_2_=5 and total TR=90ms. The AFI acquisition had twice lower resolution than GRE in each dimension. Other parameters were: BW= 285Hz/px, TE=3.23 ms, 4 avg. Two separate scans were acquired. The scans for quantitative parameter mapping were accompanied by high-resolution (0.3mm isotropic) 3D turbo spin-echo (TSE) acquisitions with many averages; these will not be discussed here. The total measurement time per brain was close to 60hrs and generally occupied the space of a week-end (Friday evening to Monday morning). Due to different sizes of the brains and the different placement of the reference probe inside the Plexiglas cylinder, the FOV varied. In order to keep within the available measurement time, the resolution was allowed to differ slightly from case to case. The values were between 0.39 x 0.39 x 0.5 mm^3^ (two brains), through 0.39 x 0.39 x 0.6 (three brains) and up to 0.52 x 0.52 x 0.6 mm^3^ (two brains). The values are listed in Table 2.

**Table 2.** Water content and relaxometry data. Bold values indicate parameters with SD below 20% over 7 brains.

| Brain nr. | H<sub>2</sub>O [%] |  |  | T<sub>1</sub> [ms] |  | T<sub>2</sub><sup>*</sup> [ms] |  | T<sub>1</sub> [ms] MT |  | T<sub>2</sub><sup>*</sup> [ms]MT |  |
| --- | --- | --- | --- | --- | --- | --- | --- | --- | --- | --- | --- |
|  | WM | GM | r | WM | GM | WM | GM | WM | GM | WM | GM |
| 1 | 70.2 | 82.0 | 1.17 | 511 | 545 | 38 | 46 | 415 | 430 | 40 | 44 |
| 2 | 75.2 | 85.9 | 1.14 | 624 | 555 | 35 | 39 | 449 | 425 | 36 | 33 |
| 3 | 74.8 | 87.0 | 1.16 | 331 | 464 | 28 | 39 | 317 | 351 | 31 | 33 |
| 4 | 74.8 | 87.5 | 1.17 | 390 | 398 | 43 | 48 | 323 | 348 | 43 | 45 |
| 5 | 73.8 | 83.0 | 1.12 | 307 | 383 | 28 | 34 | - |  | - |  |
| 6 | 73.5 | 87.6 | 1.19 | 261 | 327 | 36 | 65 | 191 | 285 | 35 | 65 |
| 7 | 68.3 | 83.2 | 1.22 | 201 | 251 | 30 | 50 | 164 | 208 | 31 | 53 |
| Mean SD | **72.9**<br>**2.6** | **85.2**<br>**2.4** | **1.17**<br>**0.03** | 375<br>148 | 418<br>112 | **34**<br>**6** | **46**<br>**10** | 310<br>115 | 341<br>85 | **36**<br>**5** | 46<br>12 |
| in vivo | 70.0 | 82.0 | 1.17 | 979 | 1538 | 52 | 60 | 615 | 972 | 52 | 58 |

In order to avoid blurring in the high-resolution scans due to the expected warming-up of the passive shims by eddy currents and accompanying field drift, an initial single-average 3D GRE scan was acquired and discarded. The remaining scans were split into reasonably short scans (less than 3 hrs each) to minimise the effect of long-term drifts in scanner components.

The *in vivo* protocol also consisted of four 3D GRE acquisitions, two without and two with a standard module MT preparation, and one AFI scan. The parameters of the 3D GRE acquisitions included: TR=50ms, α_1_ = 7° and α_2_ = 40°, 12 echoes, TE_1_ = 2.33ms, ΔTE = 2.89ms, BW=300Hz/px, matrix size = 192×156×128, voxel size 1.04×1.04×1 mm^3^, RF spoiling, assymetric echo, 6/8 partial Fourier, parallel imaging iPAT=2.

B1+ mapping with AFI was performed with same parameters used for post mortem brains but with acceleration using parallel imaging iPAT=2. The resolution was identical to that of the GRE scans. The total measurement time was 34 min.

### 2.3 Data analysis

Data processing and analysis were performed with in-house MATLAB scripts (The Mathworks, Natick, MA, USA) and SPM8 (http://www.fil.ion.ucl.ac.uk/spm).

#### 2.3.1 Registration and averaging

All separate scans were coregistered off-line. Scans acquired with the same parameter combination were averaged. Complex data were used for averaging, as described in [Oros-Peusquens et al, 2010]. However, for some of the measurements changes in the field distribution with time were noticed, leading to significant signal loss after complex averaging. In order to employ the same data evaluation procedure for all brains, we used magnitude averaging throughout.

##### Concepts

The general corrections required for accurate water content mapping have been discussed previously (Tofts, 2003; Neeb et al., 2006b; Neeb et al., 2008). The corrections relevant to the present method are briefly summarized below for consistency.

#### 2.3.2 Signal equation

The signal equation for spoiled gradient echo (FLASH) is [Haase et al. 1986]:

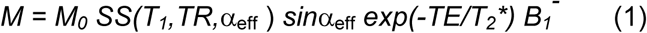

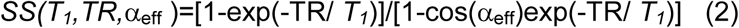

with α_eff_ = α_nom_B_1_ ^+^ where α is the effective flip angle and α is the nominal one. B_1_ ^+^ and B_1_ ^−^ are corrections reflecting deviations of the transmit and the receive field from a homogeneous distribution of the nominal flip angle and from unity, respectively. In this case, B_1_ ^+^ also includes errors in the automatic calibration procedure of the scanner. The term labelled SS is the steady-state correction to signal intensity, due to the use of a repetition time which is shorter than or comparable to T_1_.

A useful quantity can be deduced from M_0_, which is the percent tissue water content. This is often done by comparison with the signal from a reference probe of known water content. Therefore, the general form of the correction is (Neeb et al., 2006b; Neeb et al., 2008):

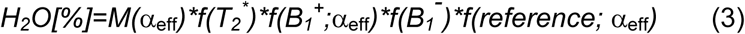

where *f(…)* are terms describing T_2_*, RF transmit, RF receive and probe calibration effects.

We note at this point that the overall correction is factorised into multiplicative terms, each depending on one parameter only. The factor *f(reference;*α) is a scaling factor which determines the correct position of the mean value of the water distribution.

The correction terms in Eq. (3) which depend on the RF field inhomogeneities are expected to have smooth and long-range variations over the brain.

#### 2.3.3 Parameter optimisation

The simulations were based on the signal equations for 3D GRE (1) and (2), which assume perfect spoiling. White noise was added to the real and imaginary parts of the signal. Receive and transmit B1 inhomogeneities were neglected in a first step. For a range of T_1_ values, the experimental parameters which deliver the best results for T_1_ and M_0_ quantification were determined, under the constraint imposed by the available measurement time that TR should not exceed 50ms for *in vivo* scanning. The maximum value TR=50ms was adopted, to allow for acquisition of echoes up to around TE=T_2_* [Oros-Peusquens et al., 2008]. The standard deviation of T_1_ and M_0_ values determined from the noisy data (SD(ΔT_1_), precision), as well as the systematic bias (mean(ΔT_1_), accuracy) in the calculated values were estimated based on 10000 simulated measurements. A T_1_ relaxation range of 200-500ms was expected for the post mortem brain (Blamire et al. 1999, McNab et al 2009, Oros-Peusquens et al. 2003); the parameters were optimised for T_1_=350ms.

For *in vivo* parameter mapping a longitudinal relaxation time of T_1_=1000ms was chosen, based on known relaxation values at 3T [e.g. Oros-Peusquens et al., 2008].

#### 2.3.4 Normalisation to known water content

Water content in the brain was deduced by comparison to the known water content (80%) of a reference probe included in the formalin-filled container. Due to the proximity of the probe to the brain and the immersion in a common, heat-conducting fluid, no correction for temperature differences between brain and probe was considered necessary. The conversion between magnetisation density and water content described here involves the assumption that the density of water is the same in the probe (bulk water) and in fixed tissue for a given temperature.

#### 2.3.5 Inhomogeneity filtering using SPM

Due to the multiplicative nature of the correction described in Eq. (3), after taking into account effects due to T_1_, T_2_* and B ^+^, both M_0_ and the correction field can be directly estimated from the corrected signal values using a probabilistic framework for segmentation (Ashburner and Friston, 2005). The unified segmentation combines image registration, tissue classification, and bias correction in a single generative model and optimizes its log-likelihood objective function.

This is a step which has become a standard procedure in brain segmentation (Ashburner and Friston, 2005), as a large number of papers in neuroscience demonstrate.

Bias field correction was performed using SPM8 (www.fil.ion.ucl.ac.uk/spm). The bias field was estimated only on brain tissue, as selected by a brain mask, but the bias field correction was calculated for the whole field-of-view, including the probe, and subsequently applied to it.

In a first step, the uncorrected high-resolution M_0_ maps obtained from post mortem brains were brought to an anatomical orientation close to axial orientation *in vivo* and resampled to a resolution of 1×1×1mm^3^.

The brain mask was generated using the brain tissue probability maps obtained from a first run of the SPM algorithm on the whole FOV: all voxels with a probability value higher than 0.25 in any kind of tissue (WM/GM/CSF) formed the brain mask. Tissue masks for each tissue type were calculated according to the following conditions:

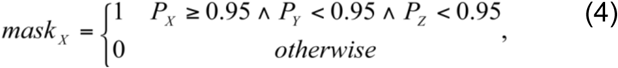

where X, Y and Z represent the three tissue types WM, GM and CSF in all possible permutations and PX is the probability of tissue X.

The bias field and the tissue masks were resampled to the original resolution of the uncorrected M_0_ map and transformed back to the original orientation. Bias field correction was applied by dividing the original intensity to the calculated inhomogeneity.

#### 2.3.6 Water maps

All corrections described above were applied to the raw data. The magnetisation density was deduced by comparing the signal intensity obtained at two different flip angles and for each echo time to the signal equations (1) and (2), T_2_* corrected by mono-exponential fitting and subsequently SPM filtered. Normalisation to the known water content of the probe was used to convert magnetisation density in gray scale values to water content in percent units.

#### 2.3.7 Magnetisation transfer

The same processing steps described above were applied to the MT-prepared scans. The quantitative information obtained was: observed equilibrium longitudinal magnetisation after magnetisation preparation M_0_ ^sat^, T_1_ after magnetisation preparation T_1_ ^sat^ , and T * after magnetisation preparation T *^sat^. By comparison with the same tissue parameters obtained without MT preparation, the following quantities were calculated:

a. Magnetisation transfer ratio

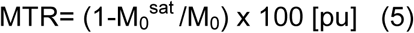 Alternatively, the ratio was calculated using the signal intensity of the first echo of the low flip angle scans

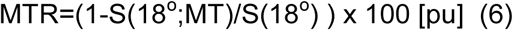
b. The pseudofirst-order rate constant for the transfer of magnetisation from the free to the bound pool [Wolff and Balaban 1989; Tofts 2003; Ropele 2000]

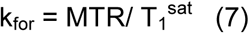 with MTR expressed as a ratio, not as percentage.
c. An approximation of the bound pool fraction

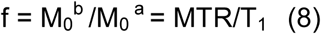 considering the saturation of the bound pool to be 1 [Lee and Dagher 1997; Tofts 2003].

#### 2.3.8 Mean values and correlation between parameters

Masks of the whole brain, WM and GM were produced for the SPM-based bias field correction, described in 2.3.5.

For each MRI-based tissue parameter determined in this study, its distribution over each tissue type and the whole brain were determined. The centroid and standard deviation were calculated.

For each combination of parameters, scatter plots were produced separately for WM, GM and the whole brain. The correlation coefficient r was calculated using the Matlab routine *corrcoef*. Thus, values of the coefficient close to 1 suggest that there is a positive linear relationship, values close to -1 suggest a negative linear relationship and values close to zero suggest no linear relationship between the two parameters.

## 3. Results

### 3.1 Parameter optimisation

Optimum parameters for 2-point relaxometry were determined as described in section 2.3.3.

For a TR=50ms and for T_1_ values around 350ms, the precision and accuracy of the method were highest for flip angles α_1_=17° and α_2_=75°. Assuming an SNR of 20 or higher in the measured data, the standard deviation for both T_1_ and M_0_ was kept below 10%. Systematic deviations of less than 5% were obtained. The accuracy and precision of the method were found to be higher by around a factor of 2 for the determination of M_0_ than for T_1_. The performance of the method improves with SNR.

Optimum flip angles of α_1_=7° and α_2_=40° and similar results regarding precision and accuracy around the optimum were obtained for T_1_ around 1000ms. This is a representative value for *in vivo* longitudinal relaxation. However, due to the broad distribution of T_1_ values in brain tissue, roughly 500 to 2000ms at 3T [Oros-Peusquens et al., 2008], larger errors are predicted for the high T_1_ values. Fig. 1 shows the optimisation results relevant to fixed tissue.

**Fig. 1.**
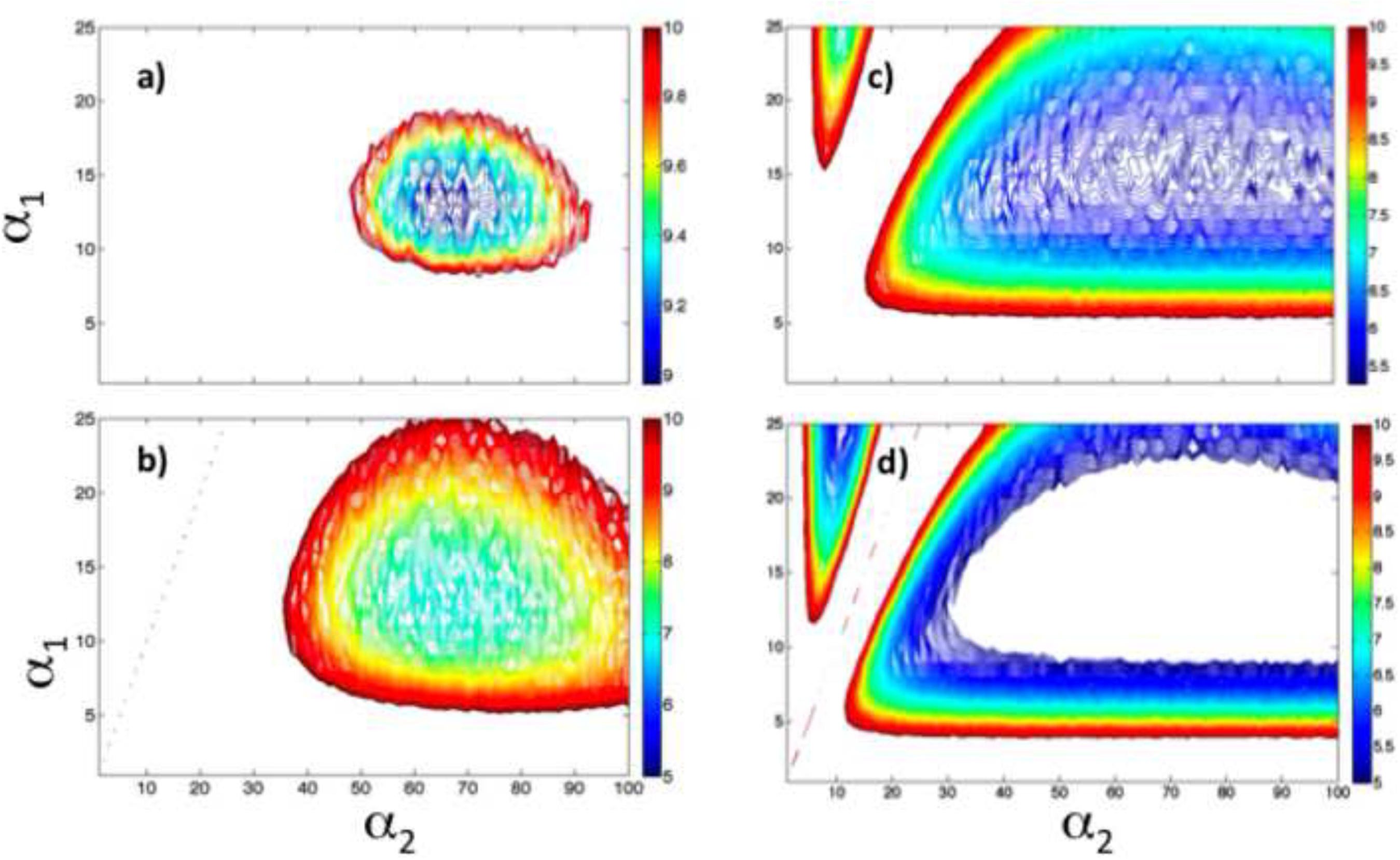
Parameter optimization for the 2-point mapping method when TR_1_=TR_2_=50ms and SNR=20. Dependence on α_1_ and α_2_ of: a) SD(T_1_); b) Δ( T_1_); c) SD(M_0_); d) Δ( M_0_). Only the combinations of parameters are shown for which a-d remain below 10%.

### 3.2 SPM bias field correction

The quality of the segmentation into WM, GM and CSF was good (Fig. 2a and b show masks for GM and WM), as subsequently determined by visual inspection. After bias field correction the water map shows excellent tissue contrast with no apparent long-range variations (Fig. 2c) and the histogram of water content distribution in the brain shows two well separated peaks corresponding to WM and GM (Fig. 2e).

**Fig. 2.**
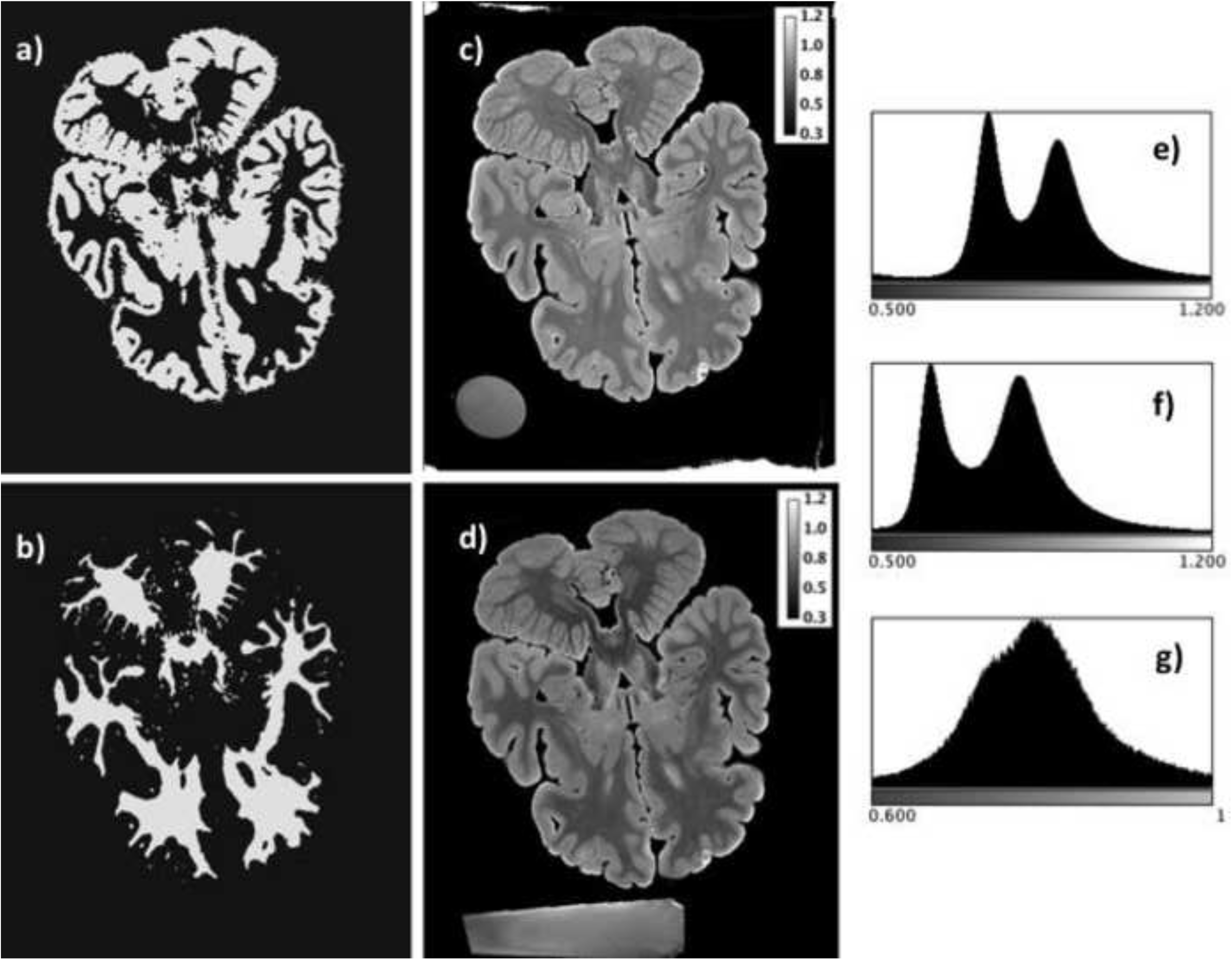
a) Mask for GM; b) mask for WM; c) water content; d) bias-field corrected map of the equilibrium magnetisation after MT; e) histogram of water content distribution in the brain; f) histogram of the equilibrium magnetisation after MT; g) histogram of „water content‟ value distribution in the probe after correction with extrapolated bias field. For normalisation to the known water content of the probe (80%), ROIs close to the brain were chosen, where the extrapolation error for the bias field is small.

The bias field correction for the M ^sat^ maps was performed independently; the results regarding segmentation and bias field correction were very similar to those obtained for M_0_ maps. The map of „water content after MT‟ reflecting the longitudinal equilibrium magnetisation in the presence of saturation, is shown in Fig. 2d and its histogram in Fig. 2f.

In most cases, the width of the intensity distribution in the water probe after correction (the bias field was extrapolated to cover the whole FOV) was still large, in the range of 20% (Fig. 2g). Therefore, restricted homogeneous regions within the probe, which were spatially closest to the brain, were used for calibration of M_0_ maps to water content.

### 3.3 Contrast

As usual in 2-point T_1_ mapping protocols, the image obtained with low flip angle is M_0_ weighted and the one obtained with the large flip angle is T_1_ weighted. Superimposed on these initial contrasts, T_2_* contrast increases with increasing echo time. The MT-prepared scans have additional MT contrast. Tissue characterisation (for example, segmentation) can be done based on the quantitative parameters, and also on the original contrasts, even if they are redundant. One of the advantages of using the original contrasts is their higher SNR compared to the quantitative maps.

We show in Fig. 3a an example of the 6+6+4+4=20 original contrast images for a representative brain and slice. Fig. 3b shows a magnified region including the thalamus, reflected in the first four principal components extracted from the 20 contrasts (normalised to the mode of the intensity distribution). To illustrate the potential for automatic segmentation of such data, we show results of k-mean clustering with an arbitrary number of 16 clusters performed either on the 20 contrasts (top) or on the first four PCAs (bottom). The figure is only meant to illustrate the potential of the method – for a proper study regarding segmentation of the thalamus or other regions based on the data from the seven brains, additional work is required [see e.g. Deoni et al. 2007].

**Fig. 3.**
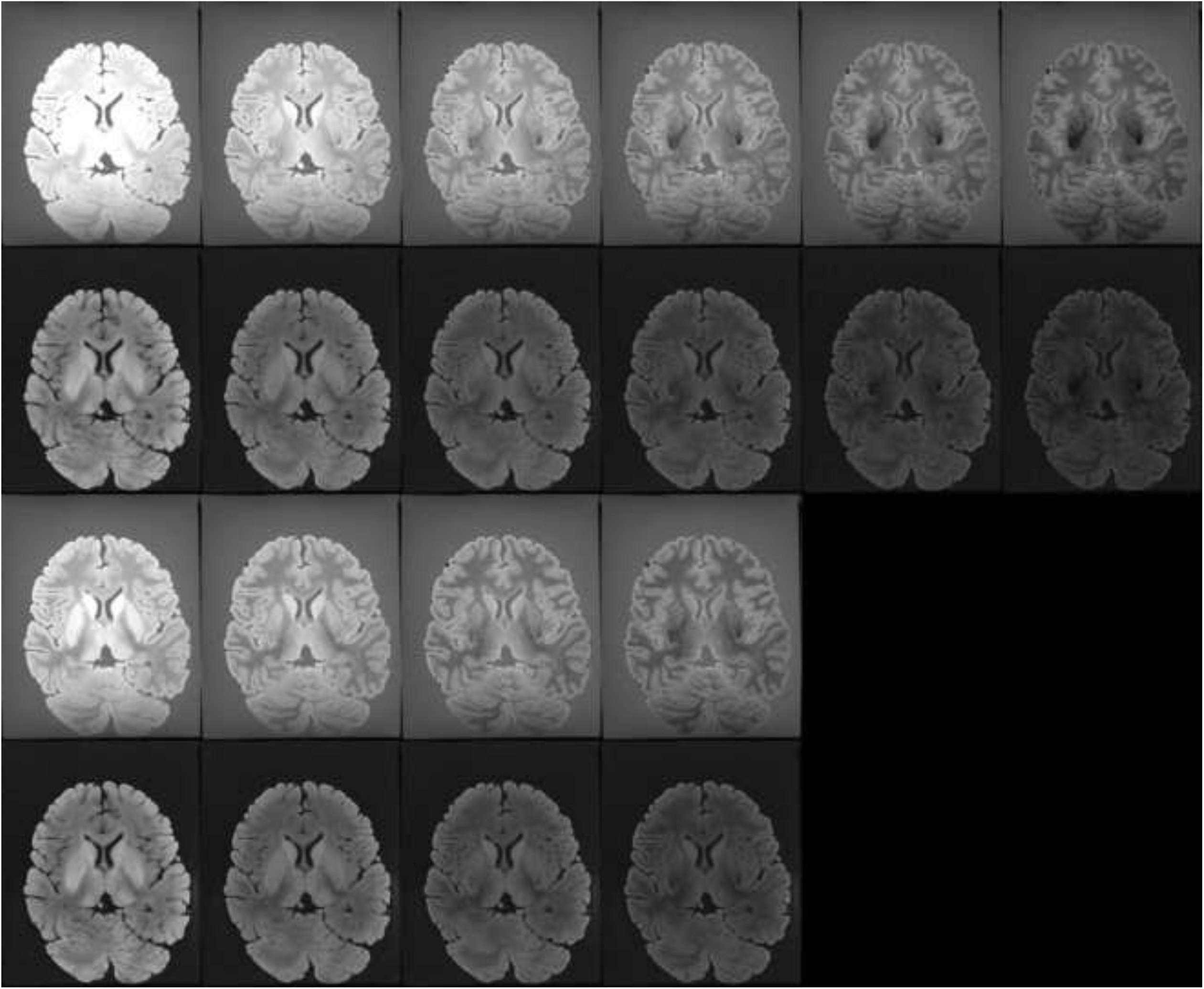
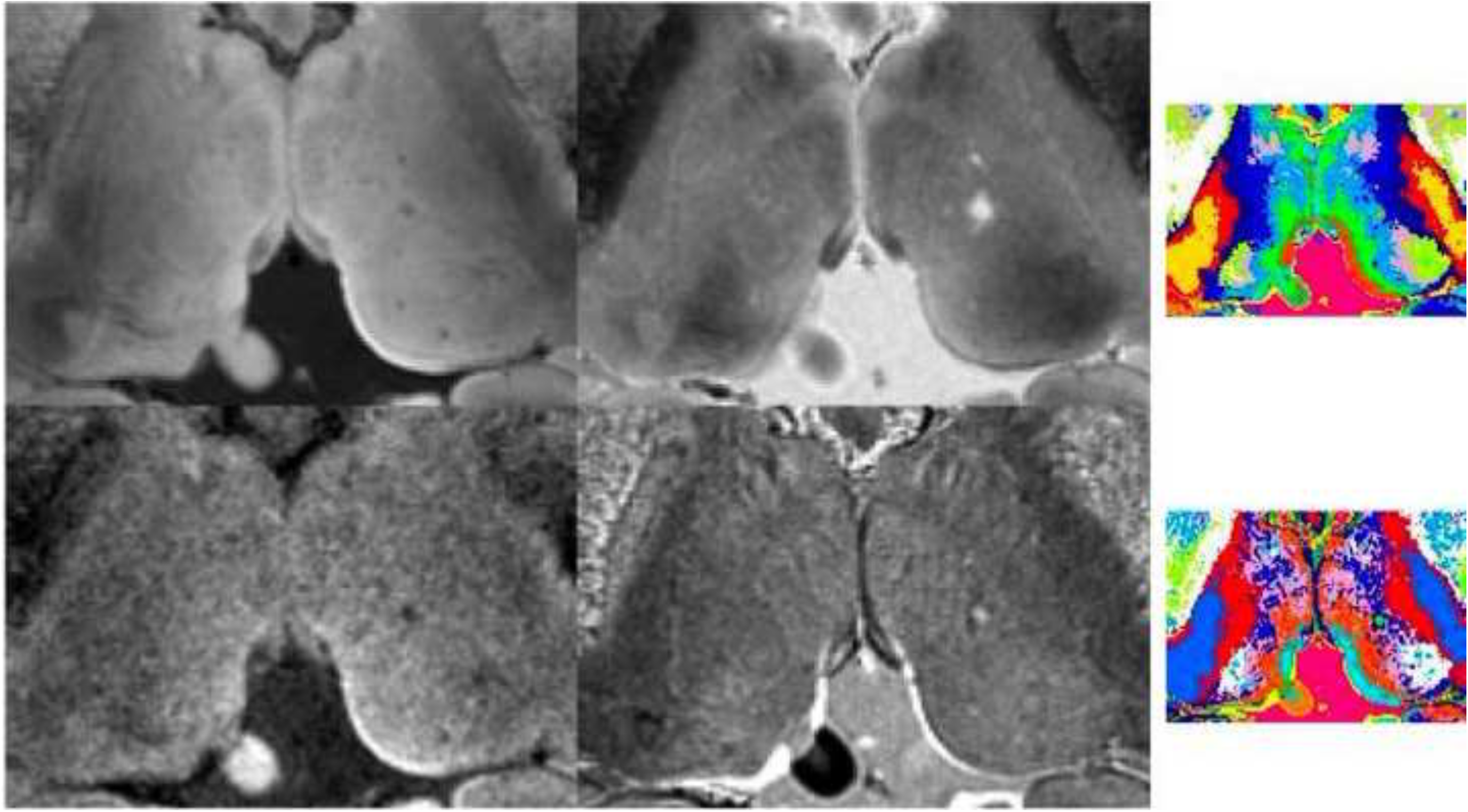
a) Multi-contrast data from which the quantitative maps are calculated. From top to bottom: 6-echo low flip angle data set; 6-echo high flip angle data set; 4-echo low flip angle MT-prepared data set; 4-echo high flip angle MT-prepared data set. b) Magnified region containing the thalamus: grey-scale images are the first four PCAs from the contrasts shown in a). Color images are k-means clustering results obtained from all contrasts (right, top) or from first four PCAs (right, bottom). An arbitrary number of 16 clusters have been used for illustration.

### 3.4 Quantitative parameter maps

The multiple contrasts reflect 6 parameters: M_0_, M_0_ ^sat^, T , T ^sat^ , T * and T *^sat^. Another three parameters were derived as described in 2.3.7: MTR, k_for_ and f. Fig. 4 shows the 9 quantitative parameters listed above, with the addition of T_1_/T_2_*, a quantity similar to the mixed contrast T_1_/T_2_ images suggested to reflect myelination *in vivo* [Glasser and van Essen 2011]. Brain 1 was chosen for representation purposes. Four maps with high contrast, M_0_ ^sat^, T /T *, f and T * are shown in three orthogonal orientations in Fig. 5 a-d, to allow the reader to appreciate the level of anatomical detail visible with very high contrast in the maps.

**Fig. 4.**
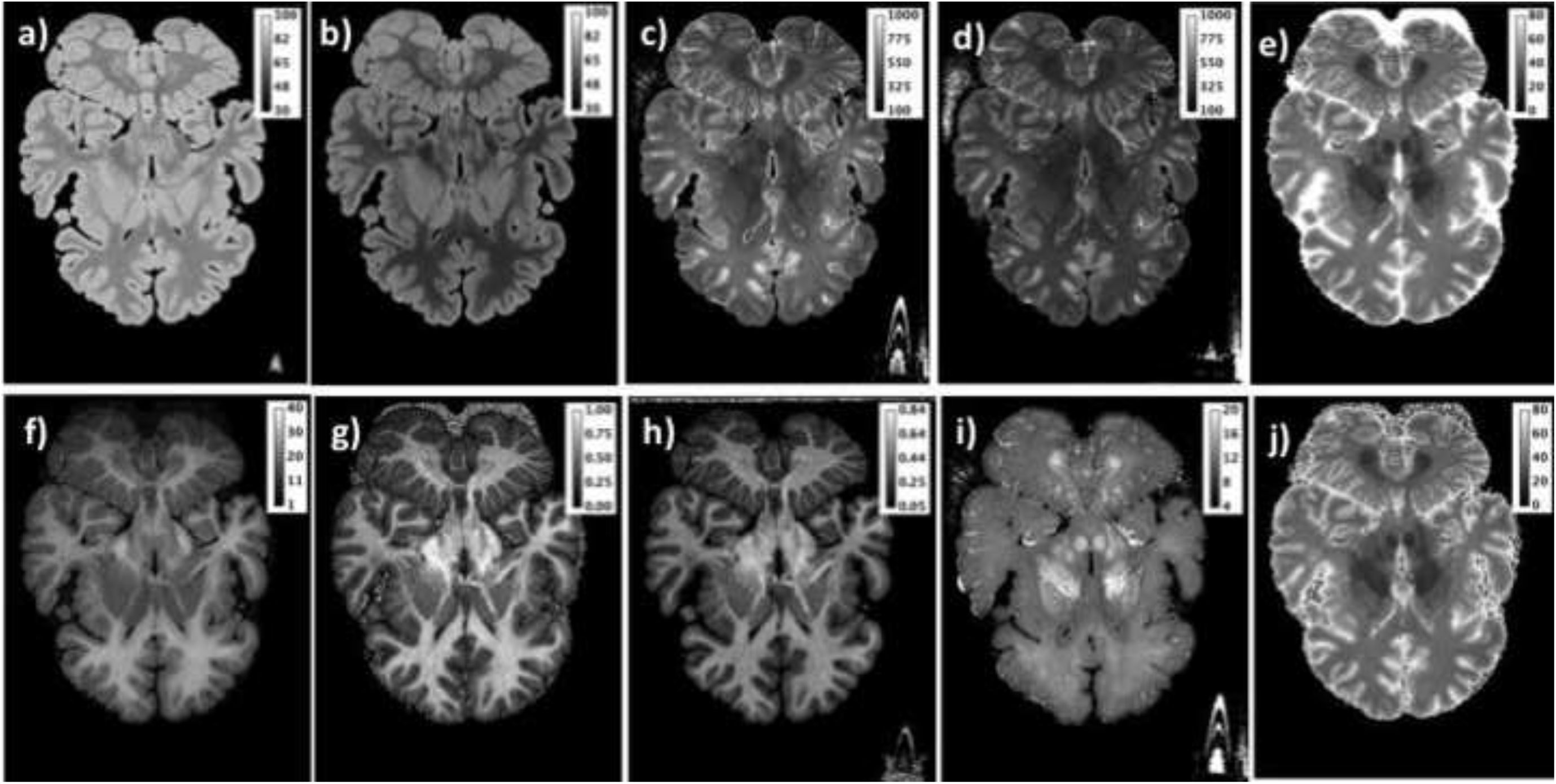
Quantitative maps of a representative post mortem brain (nr. 1): a) water content in percent units [pu]; b) bias-field corrected map of the equilibrium magnetisation after MT (M_0_^sat^, pu); c) T_1_ map [ms]; d) T_1_ after MT (T_1_^sat^) [ms]; e) T_2_* [ms]; f) MTR [pu] ; g) k_for_ [s^−1^]; h) f_bound_ [a.u.] i) T_1_/T_2_*; j) T_2_*^sat^ [ms].

**Fig. 5.**
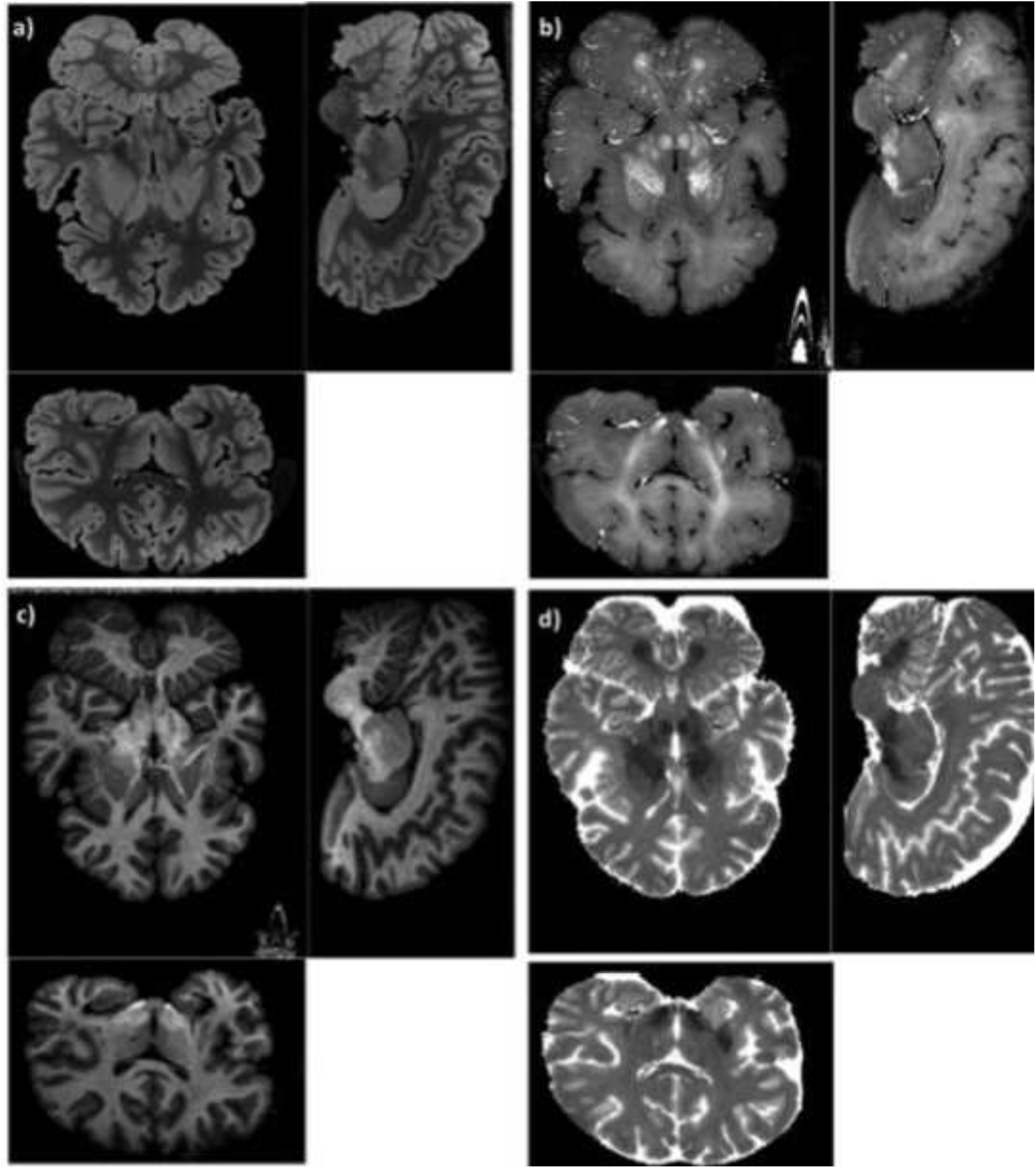
High-contrast maps shown in all three orthogonal orientations: a) M ^sat^, b) T_1_/T_2_*, c) f_bound_ and d) T_2_*.

Figure 6 shows examples of similar quantitative maps obtained *in vivo*. The selection consists of: water content (a), T_1_ (b), T_1_ ^sat^ (c), T * (d), MTR (e), k_for_ (f), bound fraction f (g) and T_2_*/ T_1_ (h). Another example of quantitative maps is given in Fig. 7 for a post mortem brain (nr. 4) showing a broad rim fixation artefact. The upper row shows maps which are insensitive to the artefact: water content (a), M_0_^sat^ (b), k_for_ (c), and the lower row shows maps in which the artefact is well visible: T_1_ (d), T_2_* (e), MTR (f).

**Fig. 6.**
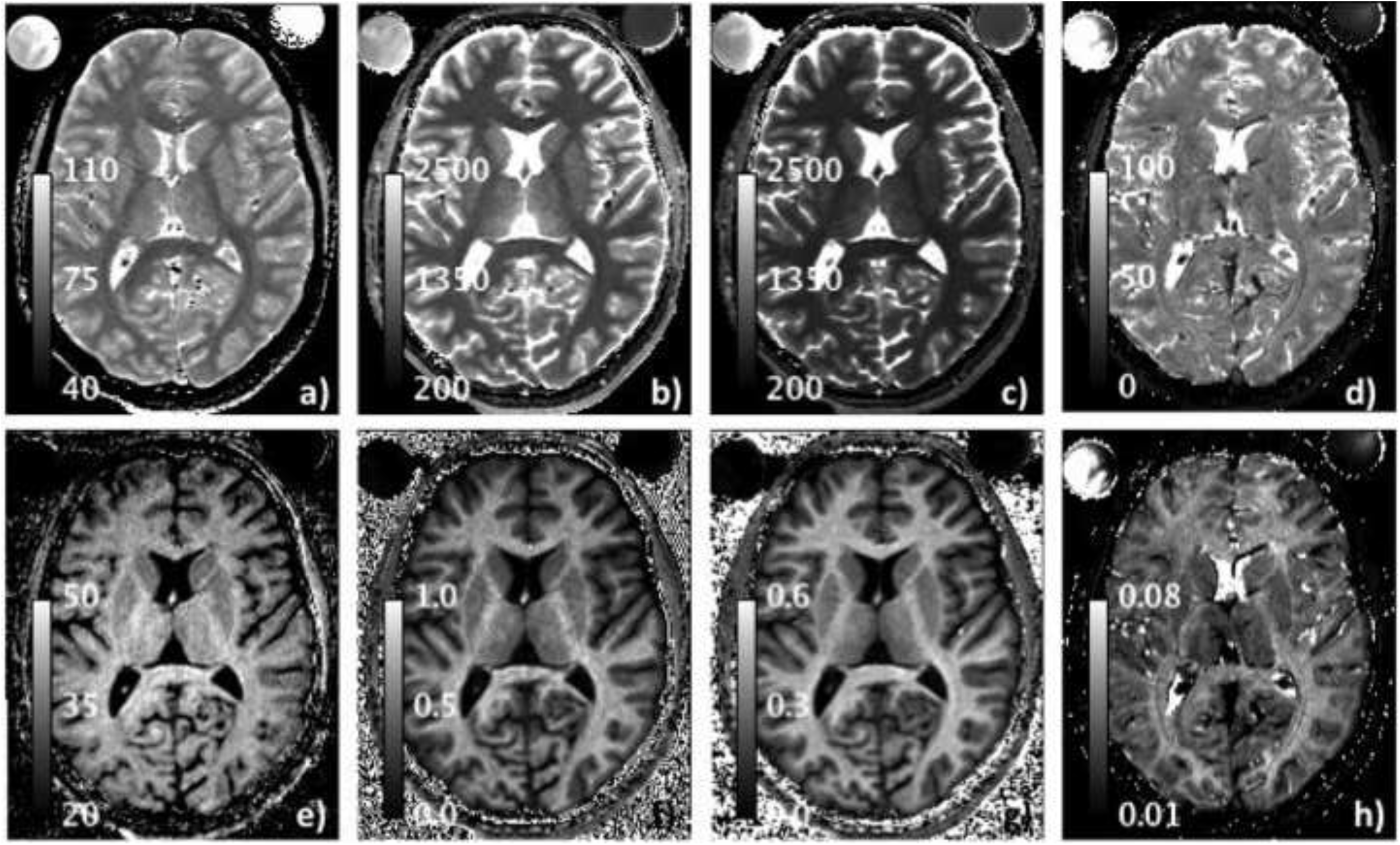
Quantitative maps *in vivo*. a) water content in percent units [pu]; b) T_1_ [ms]; c) T ^sat^ [ms]; d) T * [ms]; e) MTR [pu] ; f) k [s^−1^]; g) f [a.u.]; h) T /T *.

**Fig. 7.**
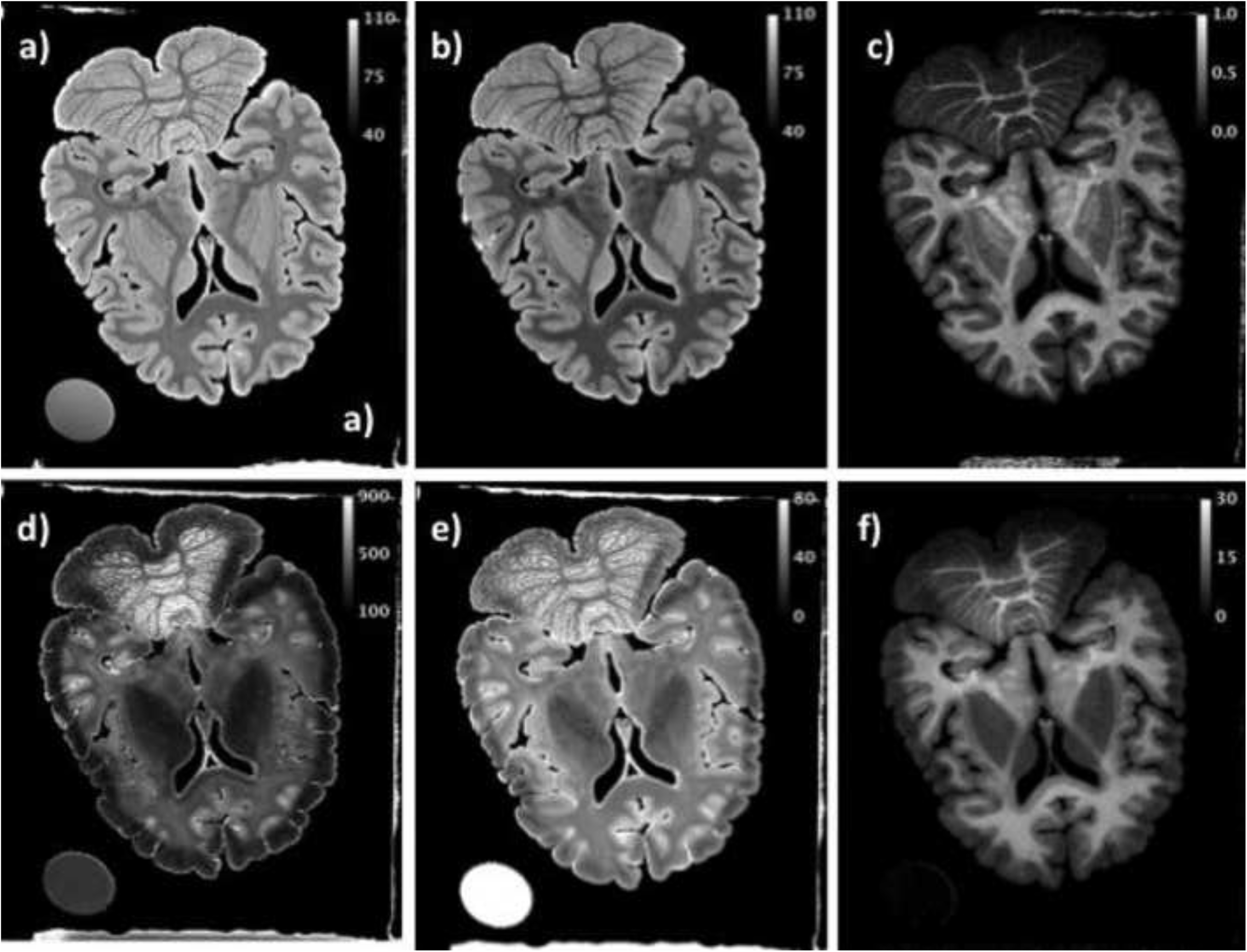
Maps of a post mortem brain (nr. 4) showing a fixation artefact. a) water content in percent units [pu]; b) bias-field corrected map of the equilibrium magnetisation after MT (M ^sat^, pu); c) g) k [s^−1^]; d) T [ms]; e) T * [ms]; f) MTR [pu].

The mean values of water content, relaxation times and MT parameters are listed in Tables 2 and 3 for the 7 fixed brains and for the *in vivo* case. Tissue masks obtained with SPM were applied to brain tissue. The values are listed separately for WM, GM and whole brain. The mean values and standard deviation over the 7 fixed brains are calculated. The values highlighted in bold face represent quantities for which the standard deviation was found to be less than 20% of the mean value when calculated over all 7 brains.

**Table 3.** Parameters characterising magnetisation transfer.

| Brain<br>nr. | MTR [%] |  | k [s<sup>-1</sup>] |  | f [a.u.] |  |
| --- | --- | --- | --- | --- | --- | --- |
|  | WM | GM | WM | GM | WM | GM |
| 1 | 24.3 | 16.1 | 0.66 | 0.37 | 0.47 | 0.30 |
| 2 | 28.8 | 16.9 | 0.67 | 0.32 | 0.43 | 0.28 |
| 3 | 21.8 | 14.7 | 0.84 | 0.37 | 0.66 | 0.33 |
| 4 | 16.1 | 9.9 | 0.51 | 0.30 | 0.42 | 0.26 |
| 5 | 12.4 | 9.2 | - |  | 0.42 | 0.25 |
| 6 | 25.5 | 18.5 | 1.37 | 0.65 | 1.01 | 0.56 |
| 7 | 22.9 | 16.6 | 1.34 | 0.69 | 1.05 | 0.59 |
| Mean | 21.7 | 14.6 | 0.90 | 0.45 | 0.64 | 0.37 |
| SD | 5.7 | 3.6 | 0.37 | 0.17 | 0.28 | 0.14 |
| in<br>vivo | 40.1 | 34.0 | 0.61 | 0.35 | 0.41 | 0.23 |

Fig. 8 shows M_0_ (a), T_1_ (b) and T_2_* (c) maps for each of the fixed brains; a slice with similar anatomy was chosen in each case. Beside a large variation in the mean whole-brain T_1_ value, also a significant variability in the WM-GM T_1_ contrast from brain to brain is observed. In contrast, the water content distribution was found to be very similar for all brains and offer excellent WM-GM contrast. Furthermore, and unlike the *in vivo* case [Oros-Peusquens et al., 2008], T_2_* values showed narrow intervals of variation per tissue class, good WM-GM contrast in most cases and rather constant values over the seven brains.

**Fig. 8.**
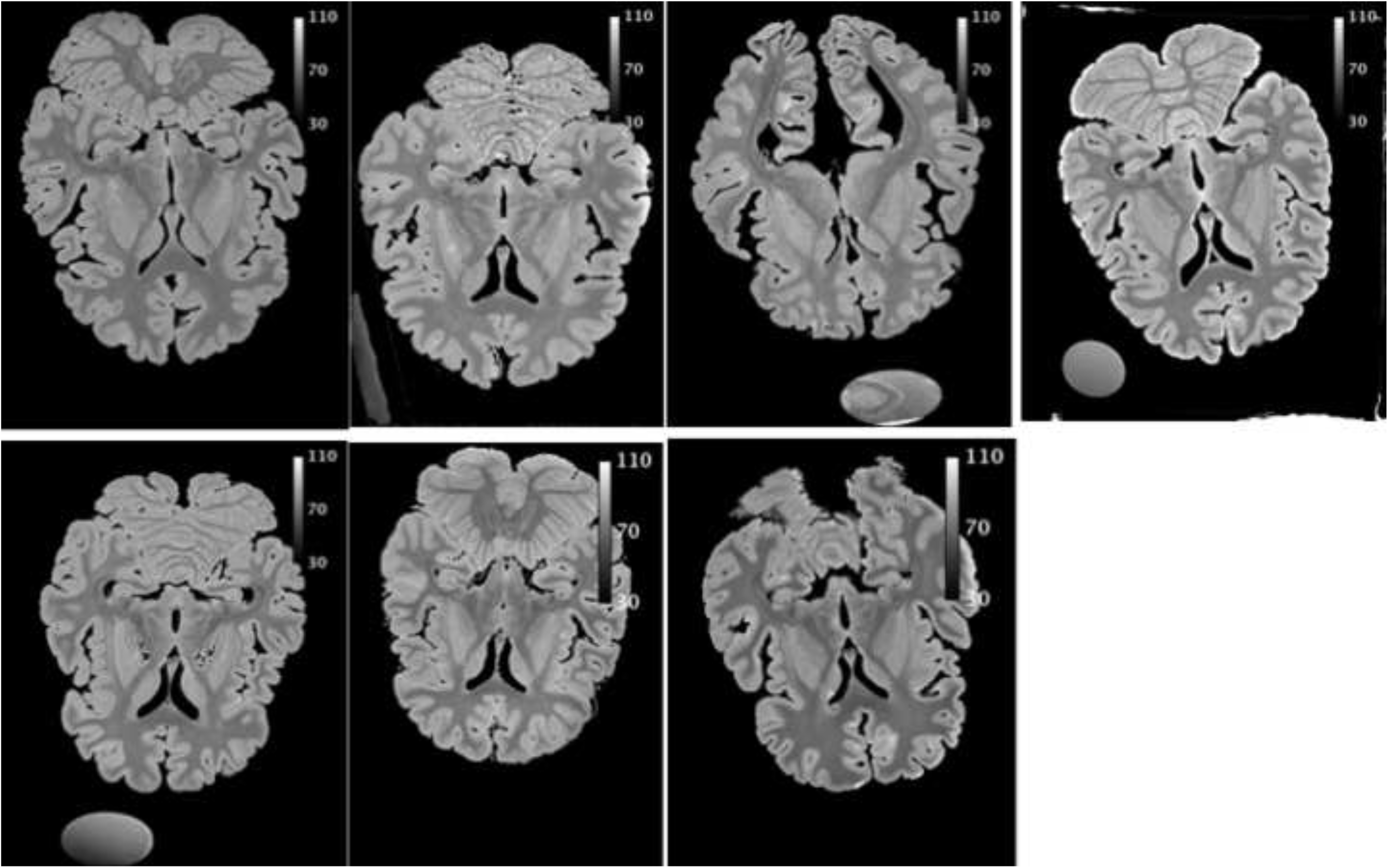
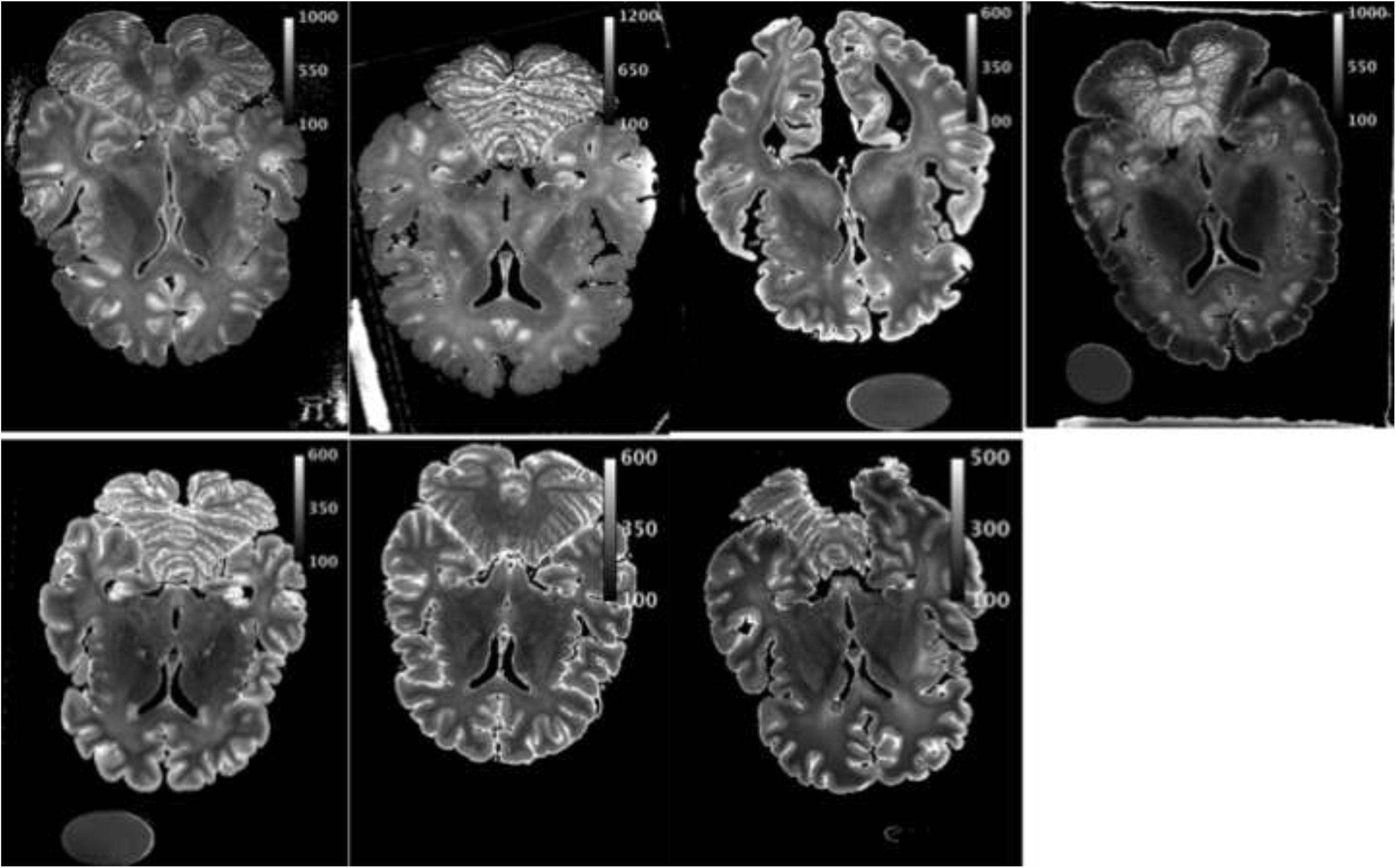
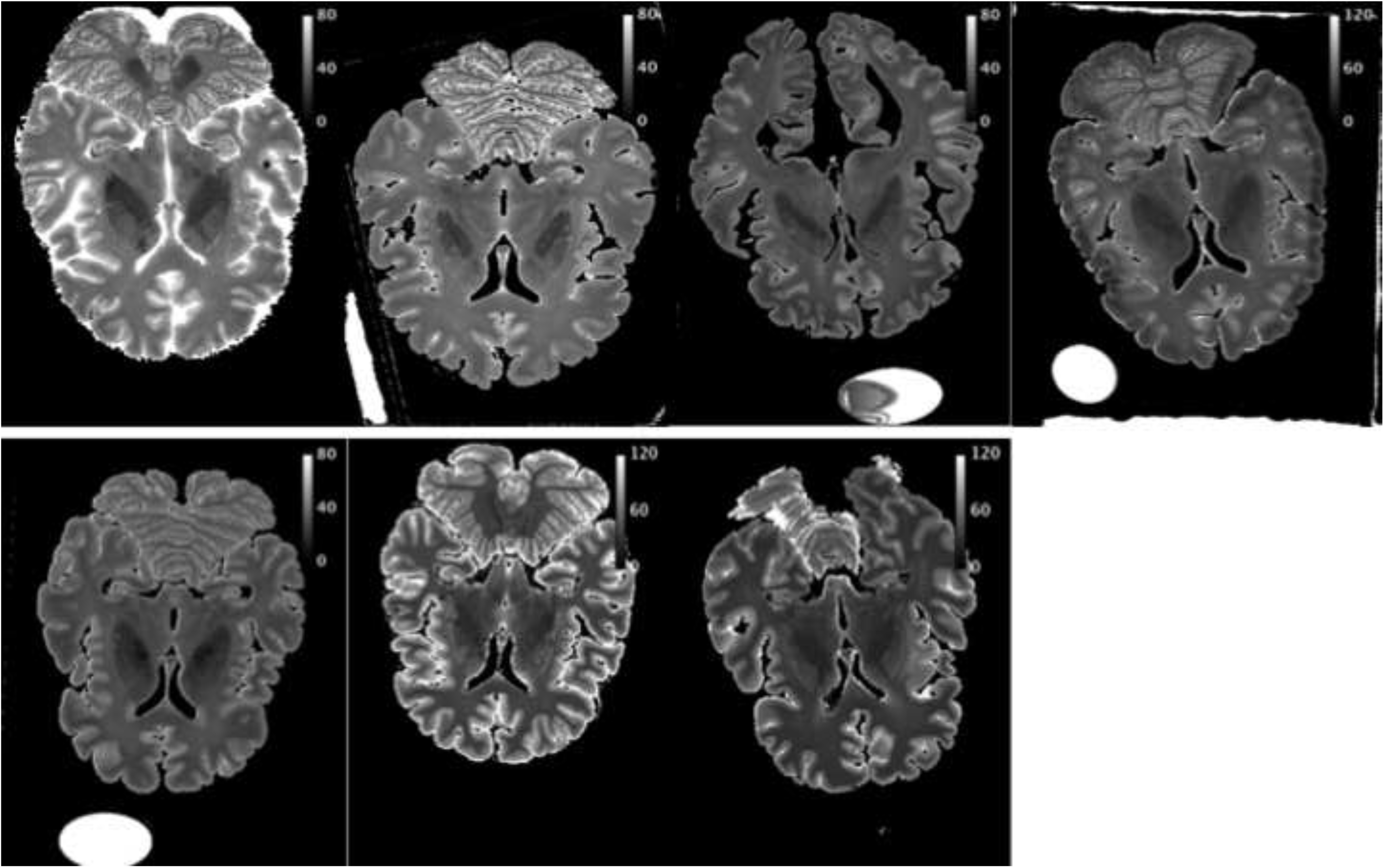
a) M_0_, b) T_1_, and c) T_2_* maps for a similar slice in each of the 7 fixed brains.

### 3.4 Correlations between quantitative parameters

Fig. 9 a-d and Tables 4 and 5 characterise the correlation between water content and different other quantities in fixed tissue. Fig. 9 includes only data for one of the fixed brains (brain nr. 1).

**Fig. 9.**
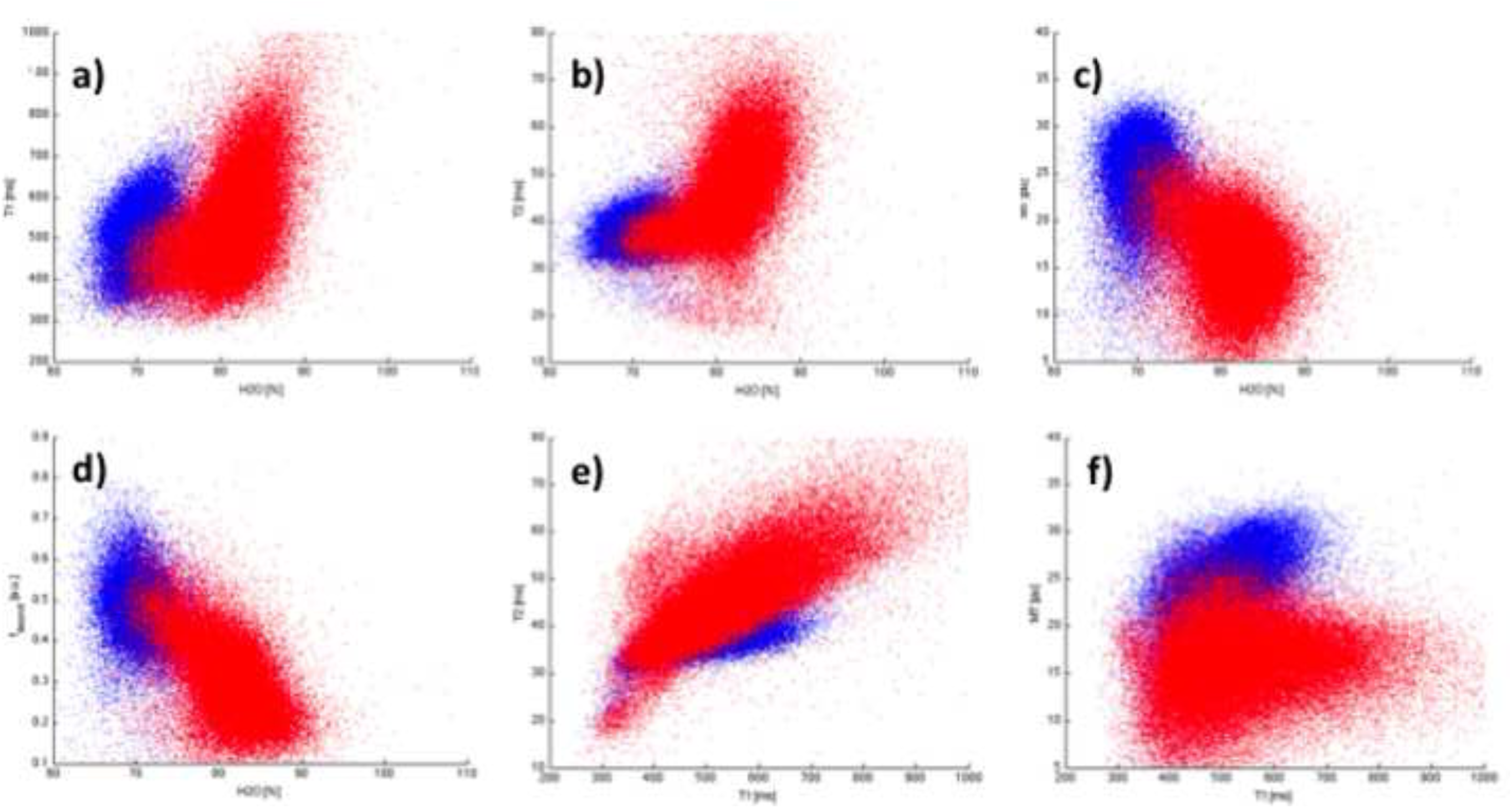
Correlation between selected quantities in fixed tissue. Blue: WM, red: GM. a) water content (H_2_O) - T_1_; b) H_2_O - T_2_*; c) H_2_O – MT; d) H_2_O – f_bound_ ; e) T_1_ -T_2_*; f) T_1_ – MT.

**Table 4.** Correlation coefficients: relation between water content and other parameters. Bold values indicate parameters with SD below 20% over 7 brains.

| Brain<br>nr. | H<sub>2</sub>O - T<sub>1</sub> |  |  | H<sub>2</sub>O - T<sub>2</sub> |  |  | H<sub>2</sub>O - MTR |  |  | H<sub>2</sub>O - k |  |  | H<sub>2</sub>O - f<sub>bound</sub> |  |  |
| --- | --- | --- | --- | --- | --- | --- | --- | --- | --- | --- | --- | --- | --- | --- | --- |
|  | W<br>M | GM | brain | WM | GM | brain | WM | GM | brain | WM | GM | brain | WM | GM | brain |
| 1 | .24 | .50 | .38 | .25 | .47 | .59 | -.23 | -.29 | -.67 | -.32 | -.50 | -.75 | -.40 | -.57 | -.77 |
| 2 | .13 | .27 | .13 | .27 | .33 | .75 | -.30 | -.26 | -.74 | -.42 | -.42 | -.70 | -.46 | -.44 | -.71 |
| 3 | .16 | .24 | .65 | .13 | .28 | .79 | .04 | .20 | -.51 | -.02* | .02* | -.75 | -.07 | .01* | -.74 |
| 4 | .20 | .26 | .18 | .33 | .28 | .39 | -.25 | -.28 | -.63 | -.39 | -.51 | -.73 | -.46 | -.57 | -.76 |
| 5 | .41 | .48 | .58 | .50 | .45 | .60 | -.53 | -.38 | -.65 | - |  |  | -.67 | -.61 | -.79 |
| 6 | .46 | .30 | .57 | .63 | .27 | .72 | -.50 | -.08 | -.63 | -.60 | -.23 | -.74 | -.64 | -.28 | -.76 |
| 7 | .33 | .65 | .57 | .48 | .54 | .78 | -.27 | -.32 | -.56 | -.52 | -.67 | -.77 | -.50 | -.69 | -.75 |
| Mean | .28 | .39 | .44 | .37 | .37 | **.66** | -.29 | -.20 | **-.63** | -.38 | -.38 | **-.74** | -.45 | -.45 | **-.75** |
| SD | .13 | .16 | .21 | .17 | .11 | **.14** | .19 | .20 | **.07** | .20 | .24 | **.02** | .20 | .24 | **.03** |
| in<br>vivo | .28 | .62 | .84 | -.12 | .12 | .34 | .09 | -.16 | -.49 | -.03 | -.31 | -.69 | -.19 | -.54 | -.82 |

**Table 5.** Correlation coefficients: relation between quantitative parameters other than water content. Bold values indicate parameters with SD below 20% over 7 brains.

| Brain<br>nr. | T<sub>1</sub> - T<sub>2</sub> |  |  | T<sub>1</sub> – MTR |  |  | T<sub>2</sub> – MTR |  |  | T<sub>1</sub> -k |  |  | T<sub>1</sub> – f_bound |  |  | k – f_bound |  |  |
| --- | --- | --- | --- | --- | --- | --- | --- | --- | --- | --- | --- | --- | --- | --- | --- | --- | --- | --- |
|  | WM | GM | brain | WM | GM | brain | WM | GM | brain | WM | GM | brain | WM | GM | brain | WM | GM | brain |
| 1 | .57 | .75 | .70 | .45 | .11 | .05 | -.01* | -.18 | -.40 | -.06 | -.37 | -.30 | -.39 | -.54 | -.46 | .89 | .93 | .95 |
| 2 | .68 | .63 | .57 | .57 | .33 | .29 | .13 | -.19 | -.37 | .02* | -.24 | -.10 | -.40 | -.44 | -.30 | .85 | .91 | .94 |
| 3 | .37 | .33 | .69 | .37 | .29 | -.26 | .10 | -.03 | -.54 | -.05 | -.36 | -.65 | -.24 | -.41 | -.68 | .92 | .97 | .98 |
| 4 | .58 | .72 | .66 | .51 | .48 | .34 | .21 | .25 | -.02 | -.23 | -.37 | -.25 | -.42 | -.51 | -.37 | .95 | .96 | .97 |
| 5 | .72 | .76 | .79 | -.05 | -.09 | -.28 | -.32 | -.31 | -.47 | - |  |  | -.62 | -.69 | -.70 | - |  |  |
| 6 | .52 | .18 | .54 | -.25 | -.30 | -.45 | -.47 | -.72 | -.56 | -.74 | -.03* | -.66 | -.56 | -.54 | -.77 | .94 | .87 | .96 |
| 7 | .42 | .60 | .55 | .39 | -.14 | .02* | -.25 | -.40 | -.57 | -.56 | -.71 | -.65 | -.73 | -.78 | -.76 | .96 | .98 | .98 |
| Mean | .55 | .57 | **.64** | .28 | .10 | -.04 | -.09 | -.23 | -.42 | -.28 | -.35 | -.44 | -.48 | -.56 | -.58 | **.92** | **.94** | **.96** |
| SD | .13 | .22 | **.09** | .31 | .28 | .30 | .26 | .30 | .19 | .31 | .22 | .25 | .17 | .13 | .20 | **.04** | **.04** | **.02** |
| in vivo | .28 | .45 | .53 | -.03 | -.35 | -.56 | -.07 | -.29 | -.39 | -.45 | -.60 | -.79 | -.78 | -.83 | -.91 | .75 | .81 | .91 |

Fig. 9e-f and Table 5 characterise correlations between parameters other than water content.

Also in this case the WM and GM masks obtained from SPM were used and the correlations were investigated separately for different tissue classes and the whole brain.

Fig. 10 reflects correlations between the same quantities as above for the *in vivo* case.

**Fig. 10.**
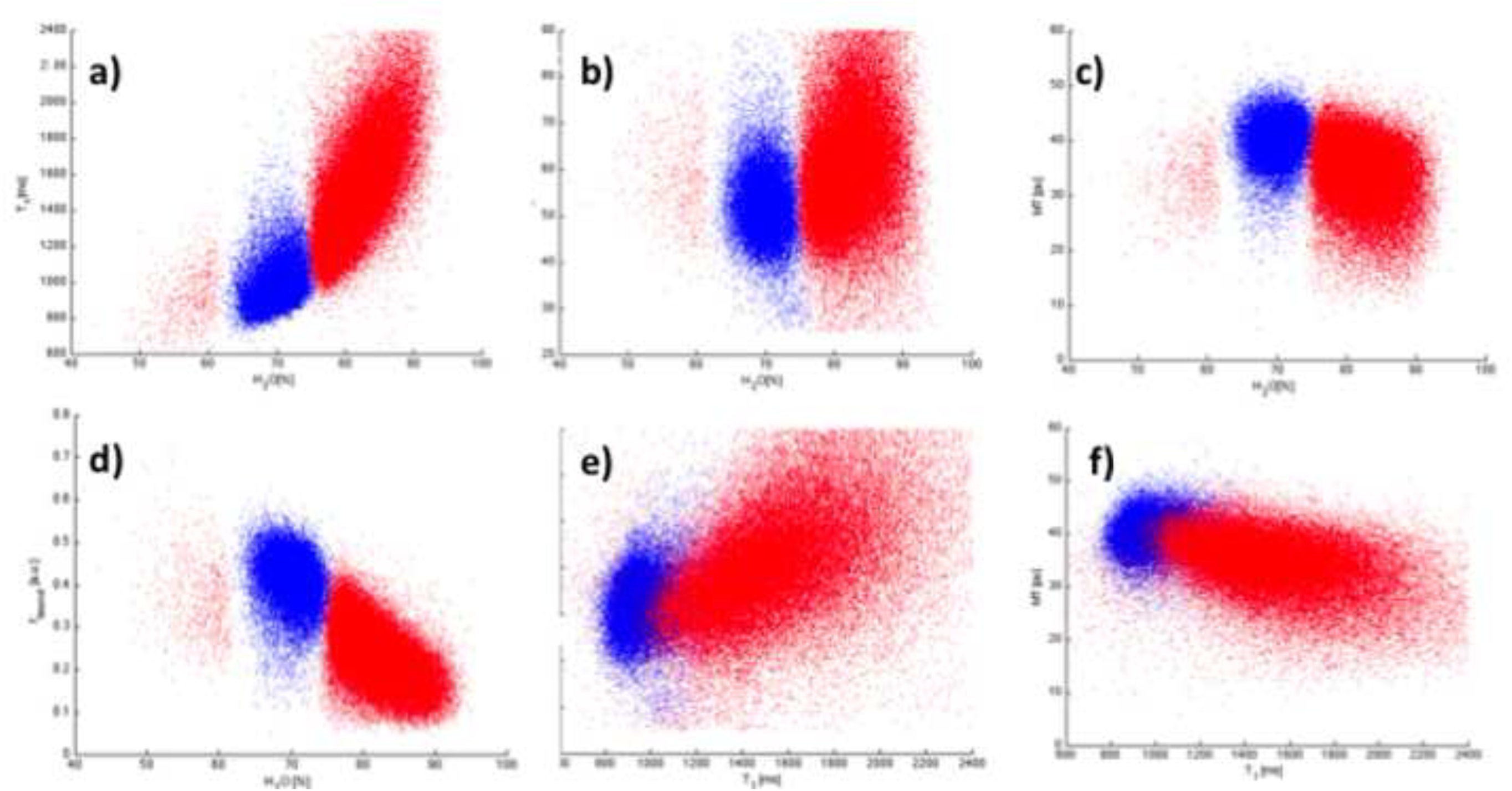
Correlation between selected quantities *in vivo*. Blue: WM, red: GM. a) water content (H_2_O) - T_1_; b) H_2_O - T_2_*; c) H_2_O – MT; d) H_2_O – f_bound_ ; e) T_1_ -T_2_*; f) T_1_ – MT.

## 4. Discussion

MR imaging performed on fixed tissue does not suffer from the same acute measurement time constraints as *in vivo* imaging. The longer available measurement time should encourage the use of quantitative methods, which are in general more time consuming that simple contrast-weighted imaging. This is due partly to the necessity to measure a number of different parameters which contribute to the determination of one quantity and partly to the fact that many quantitative methods require high SNR in the original images in order to deliver accurate results. Despite the above, and very difficult *in vivo*, high resolution is attainable even at moderate fields with quantitative imaging. One of the aims of this study was to demonstrate the feasibility of high resolution quantitative imaging at 3T for a variety of parameters. Another aim was to investigate the possibility of establishing a baseline in the values of several quantitative parameters for healthy fixed tissue and compare this with *in vivo* values. This should have impact on post-mortem examinations of pathological tissue if disturbances in one or several quantitative parameters are preserved by death and fixation.

Most of the quantitative parameters obtained in this study show large differences between fixed tissue and *in vivo* values. Furthermore, the variation of the mean values of the parameters between fixed brains is substantial. Although the variability of the quantitative values *in vivo* cannot be assessed from a single case, we can base the discussion regarding *in vivo* relaxation times and water content on data available from the literature [e.g. Oros-Peusquens et al. 2008, Neeb et al. 2006a, Neeb et al. 2006b]. We mention that M_0_, T_1_ and T_2_* - but no MT - data obtained at 3T with the same 2-point 3D method and same parameters as used for our *in vivo* case are available on a cohort of 10 volunteers. Whereas the full results, including the validation of the method on phantoms, will be discussed in detail elsewhere, we can support the discussion of the results obtained on one volunteer as being representative of a larger collective. Further data obtained on more than 30 volunteers and with different 2D methods at 3T [Oros-Peusquens et al., 2012a, Gras et al., 2011], although not included in this manuscript, contributed to assess the reliability of the quantitative results reported here for one volunteer only.

### 4.1 Water content

Water concentration is highly regulated in the healthy human brain and changes only slightly with age and sex in normal volunteers. It is affected by different pathologies such as alcoholism, haemodialysis, stroke, tumour, hepatic encephalopathy and multiple sclerosis (see [Shah et al. 2011] for a review). These changes are highly significant and thus an important parameter to be determined in the study of the disease; however, they are still only in the low percentage range. Any useful method for water content measurement must cope with this challenge of very high accuracy. This is perhaps one of the reasons why truly quantitative water measurements *in vivo* by MRI remain rare although extremely desirable. Reports of proton density changes or lack thereof with fixation can be found in the literature [Blamire et al. 1999, Pfefferbaum et al. 2004] but, to our knowledge, reports of truly quantitative water content measured by MRI did not exist for whole fixed brains before this study. By „truly quantitative‟ we mean information about water content which results in knowing the amount of water / g tissue without resorting to indirect derivations such as the one mentioned above. This is by no means a claim that relative values measured by MRI, such as relative water content of WM versus neighbouring GM, do not contain important information. Simply, one must be very careful in deriving such values to not include unwanted effects such as residual T_1_ saturation, partial volume effects and/or B_1_ ^+^ and B_1_ ^−^ inhomogeneities.

#### 4.1.1 Mapping method

Since one of our aims was to asses the existence of a baseline for quantitative values in fixed tissue, it is important to investigate the precision and accuracy of the mapping method.

The results of optimisation based on spoiled FLASH signal equation, shown in Fig. 1, indicate a sharp optimum for the precision and accuracy of T_1_ mapping (SD(ΔT_1_), mean(ΔT_1_)) with a steep increase in error outside the optimum combination of flip angles. However, please note the reduced range of variation of the errors included in Fig. 1, of only a few %. At an initial SNR of 20, the minimum bias in the measurement of T_1_ is quite high at around 6%. It does improve substantially with SNR to drop below 1% at SNR=100 (data not shown). In contrast, both the precision and accuracy of water content mapping are high and vary much more slowly outside the optimum range.

The optimum flip angles predicted by this simulation were 14.5° and 68° which are different from the **nominal** flip angles used in experiment, of 17 and 75 deg. The reason for this discrepancy was a mean deviation of the measured flip angle (by AFI, data not shown) from the nominal one of a factor 0.85. This mean value was quite constant for all brains. Since the **actual** flip angle must correspond to the optimum, the nominal flip angles were set higher.

There are known limitations of using the signal equation for perfectly spoiled FLASH for T_1_ mapping. The influence of imperfect spoiling on the mapped values was critically assessed by Preibisch and Deichmann [Preibisch 2008] for a mapping protocol corresponding to DESPOT1 [Deoni 2005]. Fortunately, we expect these effects to be very much reduced for our application. Besides using a value for the RF spoiling phase increment which reduces the variations in the derived T_1_ values (116°), we have used a TR value which is nearly 5 times longer than that used for DESPOT1 (52ms instead of 11.7ms). The longer TR used in this study was chosen based on requirements for accurate T_2_* mapping *in vivo*, but also in order to reduce the effects of imperfect spoiling on the quantitative results. Fortunately, these effects are further reduced for fixed tissue, due to the decreased relaxation times and T_1_/ T_2_ ratio.

Additional influences on the accuracy of water content mapping come from the need to use a reference value for 100% water content. *In vivo*, this is offered by CSF; for fixed brains, we have used a probe with 80% H_2_O water content and T_1_ of 320ms, very close to that expected for fixed tissue. The accuracy of parameter mapping should, therefore, be close to that for tissue. Two further possible sources of error are: (a) influence of possible signal attenuation by diffusion in imaging gradients; (b) influence of SPM bias correction. Factor (a) might be higher for post-mortem imaging than *in vivo*, due to the increased resolution and stronger gradients. On the other hand, we expect such effects to be reduced due to lower temperature and its strong influence on water diffusion. For two brains, the results obtained with the 3D method were compared to results obtained with a low resolution, very long-TR mapping method [Oros-Peusquens et al, 2012a] using lower gradient values for imaging (data not shown). No significant differences were found. Factor (b) should be considered since the bias field correction is calculated on the brain and extrapolated to the rest of the FOV. In order to minimise these effects we have used for normalisation regions in the probe which are closest to tissue. In most cases, the shortest distance between probe and tissue amounted to only a few mm.

An important step in the bias field correction is the segmentation of tissue in tissue classes. As can be seen from Fig. 2, the results are good, as assessed by visual inspection. It was important for the success of the procedure to reorient the acquired images to correspond to an axial orientation close to what would be acquired *in vivo*. Furthermore, the resolution had to be modified to 1×1×1mm^3^ and the bias field correction was initially calculated at this resolution. The inhomogeneity correction reduced the spread in grey-scale values also at probe location.

Since the SPM-based segmentation uses low-resolution prior information regarding distribution of WM, GM and CSF, we suggest that segmentation at the acquired high resolution for the post mortem brains would be better achieved from e.g. clustering procedures using multicontrast information.

#### 4.1.2 Water content maps

The distribution of water content displays very good contrast between WM and GM, as seen in Fig. 2. Moreover, this is valid for all studied brains (Fig. 8) and a remarkable constancy of the mean values over WM and GM was found for the 7 brains (Table 2). Thus, a mean value of 72.9% (p.u.) was found for WM, with a standard deviation of 2.6%(p.u.) over the 7 brains. For the GM, the values were 85.2% (p.u.) mean value, with standard deviation of 2.4% (p.u.). For comparison, the values found *in vivo* on a cohort of 10 volunteers - with the same method and parameters as reported here for the single volunteer - were 69.2% (1.9%) for WM and 81.2% (1.2%) for GM. The water content of fixed tissue is thus significantly higher (P<0.005 for both WM and GM) in fixed tissue than *in vivo*. The narrow distribution of *in vivo* values is well documented and reproducible with several methods for water content mapping [Oros-Peusquens et al. ISMRM 2011, Oros-Peusquens et al. ISMRM 2012a, Gras et al ISMRM 2011]. The high regulation of water content can be attributed to mechanisms involved in normal functioning of the brain [Pollock and Arieff 1980], which is, however, not maintained post mortem and after fixation. The constancy of water content in fixed tissue is particularly interesting given the large variation noticed for other MR parameters, most notably T_1_ relaxation.

We mention that the collective studied *in vivo* consisted of ten 26 year old male volunteers, whereas the mean age of the brain donors was 67 years with 4 female and 3 male brains. A better matched comparison of water content values should be pursued; however, the report by Neeb et al. [Neeb et al. 2006a] points toward a decrease of water content in the GM with age *in vivo*, and constancy of water content in the WM. This would further increase the significance of the heightened water content observed in fixed tissue GM.

The water content of GM cited here is the mean value of the distribution. Several regions, such as deep grey matter structures, which have higher than average water content also *in vivo*, were found to have water content above 90%.

Our results compare well with values from literature obtained from invasive as well as MRI-based measurements of water content. Fatouros and Marmarou reported a water content value for posterior white matter of 69.6% and a gray matter water content in the head of the caudate nucleus of 80.3% based on a correlation analysis between T_1_ and water content (Fatouros and Marmarou, 1999). Neeb et al [Neeb et al. 2006a], using a 2D MRI-based mapping method at 1.5T, report values of 70.9(1.1)% for white matter and 81.2(1.2)% for grey matter. Results from invasive water content measurement in biopsy samples range from 68.7% to 71.6% for white and from 80.5% to 84.6% for grey matter (Fatouros and Marmarou, 1999; Bell et al., 1987). Furthermore, for dog brain and fresh tissue, values of 84% (deep grey matter) and 74% (WM) were found (Schepps and Foster 1980), and for cat fresh brain tissue values of 71.4% (WM) and 80.8% (GM) were reported [Bakay et al Exp Brain Res 1975]. These values are in very good agreement with the average water content of 69.2(1.9)% for white matter and 81.2(1.2)% for grey matter determined here.

For fixed tissue the data are scarce; indeed, to the best of our knowledge, no MRI report of quantitative water content in fixed brain tissue existed before this study. Qualitative and partly diverging affirmations can be found: formalin-fixed tissue should exhibit slight reduction of the water content [Tovi and Ericsson 1992]; fixation modifies the water content, thus changing the proton density of the tissue and consequently the intensity of the MRI signal [Grinberg et al. 2008]; dehydration of tissue occurring post mortem [Schmierer et al. 2008]; reduction in „PD signals‟ (in grayscale units) by 20-30% over app. 30 days of fixation reported by Yong-Hing et al [Yong-Hing et al., 2005]; fixed tissues suffer reduced proton density [Miller et al 2011]; ratio of grey/white matter proton density in fixed brains is similar to *in vivo* [Pfefferbaum et al. 2004].

More quantitatively, an early CT study reports approximately unchanged water content in the brain with formalin fixation [Torack et al. 1976].

A few invasive water content measurements in fixed tissue other than brain are available. A study on rat liver and spleen with invasive water content measurement [Thickman et al , Radiology 1983] showed decreased T_1_ and T_2_ times after fixation but unchanged water content.

Fixed and living human muscle samples were investigated by invasive methods in an interesting study by Ward and Lieber (Ward and Lieber, 2005). Living muscle samples were found to have water content of 77%±2% and compared to tissue immersion-fixed in 10% buffered formalin or perfusion-fixed in 37% formalin. The tissue was either fully fixed or rehydrated by being placed in 0.2 M phosphate-buffered saline (PBS) for 0, 6, 12, 18, 24, or 30 h. Although the mean value of water content for 4% formalin fixed tissue was always slightly higher than that of fresh tissue (Fig. 2 in [Ward and Lieber 2005]), post hoc testing demonstrated that the higher water content was only significant after 30 h of hydration (81% vs. 77%, respectively). In 37% samples, water content was significantly lower at 68% after fixation but also recovered the fresh tissue value over the hydration time of 24h. Given, however, the fact that 4% formalin has 92% water and 3% methyl alcohol and 37% formalin has only 42% water and 15% methyl alcohol, the differences in water content between 4% and 37% fixed tissue are rather small. The water content of tissue obviously does not equilibrate with that of the fixative solution. Very interestingly, significantly higher muscle density was found after fixation in 4% formalin than in 37% formalin-fixed and fresh muscle. The average muscle density for 4% samples was 1.112±0.006 g/cm^3^ whereas 37% samples had a density of 1.055±0.006 g/cm^3^ and the fresh tissue value (from a different study) was 1.0597 g/cm^3^. The difference in density between 4% formalin fixed muscle and fresh tissue was significant, amounted to 4.9% and did not change after (washing out of free formaldehyde by) rehydration in PBS. An increase in density after completion of fixation was reported recently for mouse brains by Weisbecker [Weisbecker 2012]. Indeed, whereas the weight of the brains return to the original value after around 200 days of fixation, the volume was found to converge to a value smaller than the original one by 9-15%. In this context it is important to point out that the present MRI-based water mapping method actually maps MR-visible proton density in tissue. The conversion to water content values tacitly assumes that water density is the same in tissue and the solution contained in the probe. According to the present study, the difference between (MR-visible) proton density in 4% formalin-fixed and living brain tissue amounts to approximately 5% in GM and 4% in WM. These differences are already corrected for temperature dependence of the density of bulk water via normalisation to the probes.

### 4.1 Changes of tissue with fixation

Despite the widespread use of fixed tissue, the mechanism of formaldehyde fixation and its effects on the properties of tissue do not seem to be fully understood. Several aspects were reviewed in an excellent paper by Fox et al [Fox et al. 1985]. Formaldehyde (CH_2_O) dissolved in water becomes hydrated and forms methylene glycol (CH_2_(OH)_2_) ; at equilibrium, the concentration of methylene glycol is much higher than that of formaldehyde. When tissue is immersed in formalin, it is penetrated rapidly by methylene glycol and the small amount of formaldehyde present in solution, both having low molecular weight and high diffusivity. The time scale of the fixation of tissue depends on the formaldehyde being consumed after forming bonds with different tissue components and more formaldehyde forming from dissociation of methylene glycol. Thus, the equilibrium between (carbonyl) formaldehyde and methylene glycol explains why formaldehyde penetrates tissue rapidly - as methylene glycol - and fixes slowly - as carbonyl formaldehyde [Fox et al. 1985].

A further peculiar characteristic of formaldehyde fixation is vesiculation of cell membranes. Membrane changes were observed to occur in most cells exposed to formaldehyde, within a short time (20-30min) [Fox et al. 1985]. Large vesicles formed on membranes and were found to contain cytoplasmic substances.

Another unexpected effect of formalin fixation is that it causes the extracellular space in brain tissue to shrink. This is a very pronounced effect, the reduction going from around 20% in intact fresh tissue to around 5% in formalin-fixed tissue and has puzzled the electron microscopy community in its early decades [e.g. Torack 1965, Cragg 1979].

Aldehyde fixation involves chemical changes in proteins, and cell membranes as well as cytoplasmic and nuclear proteins appear to be affected [Torack 1965]. Alteration of membrane protein may easily result in a change of membrane permeability. Any effect upon the structure of cytoplasmic or nucleoplasmic protein could increase their affinity for water and by this means predipose to swelling, thus explaining the appearance of formaldehyde-fixed tissue under electron microscopy.

### 4.2 Changes of tissue MR parameters with fixation

All brains used in this study appeared at MRI to be completely fixed. Indeed, a maximum interval of 14.8 weeks for full fixation was estimated by Yong-Hing et al [Yong-Hing et al. 2005] and this was exceeded by the majority of the brains used here. Two brains showed a „fixation artefact‟ rim, which will be discussed in connection with MT results.

The observed reduction in the T_1_ and T_2_* relaxation times is in agreement with all previous reports and has been noticed very early on [Thickman 1983, Tovi and Ericsson 1992, Blamire et al 1999]. The reduction effect on T_1_ values is both very variable and very strong. On average, the values drop to one-third of the *in vivo* value and so reflect a much more pronounced effect than the reduction of only 21% reported on rat brain slices by Shepherd et al [Shepherd et al 2009]. This might be related to the duration of penetration and fixation, which were longer for the human brains due to their substantial larger size. The substantial difference in the field strength used for MRI (17.6 vs 3T) might also account for part of the effect, since it has been observed that the field dependence of living ([Rooney et al. 2007], [Oros-Peusquens et al. 2008]) and fixed tissue ([Fischer et al. 1990]) is different. In contrast a strong shortening of T_2_ values, of 81% was reported by Shepherd et al [Shepherd et al 2009] for rat brain slices, which does not seem to be met in human fixed whole brains.

We mention that most previous studies have reported changes in T_2_ whereas we have measured T_2_*. However, the shortening of 15-20 ms found in the present study in T_2_* values agrees well with the changes in T_2_ reported on human brains and also at 3T by Pfefferbaum et al. [Pfefferbaum et al., 2004].

Besides water compartmentalisation and fixative action, factors such as the distribution of paramagnetic materials and tissue inhomogeneity contribute to determine values of T_2_* and could produce variations different from those of T_2_. This was observed, for example, in a study of the effect of soaking of fixed tissue in Gd-DTPA [D‟Arceuil et al. 2007] In this context, it is interesting to notice that the vasculature, which is extremely visible in e.g in situ MRI due to the deoxygenation of blood after death, is not visible anymore in fixed tissue scans. This was observed at 8T by Dashner et al [Dashner et al, 2003] and it was also a constant finding in our MRI examinations. The explanation is most probably not the replacement of blood with formalin, as suggested by Dashner et al., since we have measured immersion-fixed brains. More likely, the source is a change in MR contrast of the blood caused by the reaction of formaldehyde with heme-linked groups [Guthe 1959] and the consequent increase in their oxygenation [Guthe 1954]). In contrast, reports of visibility of microbleeds in fixed tissue exist [Dichgans et al., 2002], also confirmed by our own observations in several fixed brains of patients (data not included in this report). In that case, the iron seems to be contained in hemosiderin [Dichgans et al., 2002].

It has been reported in a study of fixed and fresh rat brain tissue by LA-ICP-MS [Oros-Peusquens et al., 2012b] that the distribution of Fe, an element known to strongly influence R2* and R2‟=R2*-R2, does not change with tissue fixation. Blood vessels were found to still have a high iron content after fixation. The chemical state of Fe in fixed tissue was also reported to remain unchanged by Hopp et al [Hopp et al. 2010]. Several other elements, however, such as Mn, Na and K, quite radically change distribution between fresh and fixed tissue [Oros-Peusquens et al., 2012b], but since their concentration decreases, the expected effect would be a lengthening, not a shortening of the relaxation times. In the same study, the myelin content, as reflected by the distribution of phospholipids, was found to be practically unchanged. Similarly, the MRI-measured myelin water fraction was found to remain constant ex-vivo and after fixation [Moore et al., 2000]. Therefore, the observed changes in the relaxation times with fixation are not likely to be assigned to changes in either single-element distribution or myelin properties.

The reason for the relative constancy of water content and T2* values in all brains (Table 2) and the large variability of most other parameters, especially T_1_ (Table 2, Table 3) is unclear. Water is found mostly in the intracellular space, since the extracellular space shrinks to about 5% of the total [Torack 1965, Cragg 1979]. The fact that values of water content are high and comparable to results obtained invasively, suggest that most of the water is MR-visible. The reversibility of T_2_ – but not T_1_ - shortening with rehydration [Thelwall 2006, d‟Arcueil 2007], when free formaldehyde is washed out from the extracellular space, suggests that – at least after rehydration - T_2_ is determined to a large extent by a small extracellular contribution. The extent is surprising, given that the majority of water is still in the intracellular space (water content and intra/extra cellular fractions do not change with rehydration) and that T_2_ values after rehydration can even become longer that those of fresh tissue. T_1_ values are hardly influenced by rehydration, which might be a reflection of the fact that formaldehyde does not change much the T_1_ of solutions [Bossart et al 1999]. Consequently, the washing out of free formaldehyde has little influence on the T_1_ of extracellular space, but a large influence on its T_2_ [Bossart et al 1999]. Membrane permeability and exchange across membrane are reported by Shepherd et al [Shepherd et al. 2009] to be drastically increased by formalin fixation. The effect is not fully reversed by fixative wash out [Purea and Webb 2006].

A full description of this complex situation should include the intra- and extra-cellular water pools, exchange across cell membrane, exchange with the superficial water and relaxivities in the different water pools. These are a large number of parameters which cannot be directly estimated from the quantitative parameters reported here.

The pronounced tissue variability seems to be specific of human brains and not met in studies on either brain slices [Shepherd 2009, Shepherd 2009a] or sacrificed animals [d‟Arcueil 2007]. It is clear from the data contained in Table 1 and Table 2 that mean T_1_ values for WM and GM do not correlate with the post mortem interval (PMI), which is different from results reported on rat brain slices [Shepherd 2009a]. The interval between death and fixation is not the only factor influencing the properties of tissue. The best marker found for tissue quality in a recent study was RNA integrity number (Stan et al. Brain Res 2006). Factors influencing properties of tissue were: agonal factors (coma, hypoxia, hyperpyrexia at the time of death) and post-mortem interval [Stan et al., 2006]. Post-mortem degradation of RNA was not, in general, a good predictor of PMI [Sampaio-Silva et al, 2013].

It might be assumed that the uncontrolled pre- and post mortem factors influencing tissue quality account for the variability of tissue properties; in any case, these factors appear to have a larger effect on T_1_ than on T_2_* and M_0_.

### 4.3 MT parameters

The magnetisation transfer ratio (MTR) is reduced in fixed tissue, compared to the *in vivo* case (Table 3). A possible reason for this is the change in T_1_/ T_2_ (from 19 *in vivo* to 11 post mortem), thus reducing the influence of the direct effect [Henkelman et al. 2001] and the total MTR. In addition, MTR tends to reflect the product between exchange rate and T_1_ [Henkelman et al. 2001], which also decreases in post mortem tissue (∼300 compared to ∼600 *in vivo*, as obtained from mean values listed in Table 3). It is known that fixed tissue might have a different pH than *in vivo* [Ploeger et al. 1993]. From Fig. 7, the effect of the „fixation artefact‟ on T_1_ (Fig. 7d), T_2_* (7e) and MT (7f) in fixed tissue seems appreciable. The pH of the formalin solution containing the two brains with notable fixation artefacts was found to be around 5, whereas normal buffered solution has physiological pH. It can be inferred that the fixation rim has lower pH than the rest of the tissue. Interestingly, the effect of pH on T_1_ is larger than on either T_2_* or MT; k_for_ (Fig. 7c) remains unchanged. This suggests a large role played by proton exchange on T_1_, which affects T_2_* to a lesser extent. At least partly, the effect on MT might result from the combination of unaffected k_for_ and reduced T_1_ [Henkelman et al. 2001]. By affecting exchange rates and diffusion, temperature can also be expected to have a significant influence on relaxation and MT properties. Indeed, very pronounced effects of temperature were observed on T_1_ relaxation of erythrocyte ghosts [Thelwall et al 2006], with a factor 2 lower relaxation rate at 10C than at 37C. Whereas the erythrocyte ghosts model is most probably not fully representative of whole fixed post mortem brains, the point remains that care must be taken in comparing properties of fixed tissue measured at different temperatures. We have not controlled the temperature inside the scanner bore or at the sample. However, the air conditioning of the scanner room ensured constant temperature (22C) and humidity, as measured outside the bore, and given the large bore dimensions and good air circulation it might be assumed that temperature did not vary much over the approximately 60 hrs of measurement time.

We mention that the exchange rate k_for_ measured here *in vivo* is reasonably similar to that determined by Ropele et al [Ropele et al. 2000] at 1.5T using an approach similar to the present one (0.9 s^−1^ for frontal WM at 500deg flip angle). Regarding the direct comparison of values for fixed tissue to *in vivo* ones, a decrease in NAWM MTR was also reported after fixation for MS brain slices by Schmierer et al. (Schmierer et al., 2008).

We would like to stress that the model used here to extract the pseudo first order transfer rate k_for_ and the bound proton fraction f (eqs. 7 and 8) involves approximations which do not fully hold. Thus, as shown by Edzes and Samulski [Edzes and Samulski 1977] k_for_ would be the first-order rate constant for the exchange provided the macromolecular spins were kept fully saturated and provided there was no direct effect on the liquid spins. These assumptions are not fulfilled for the combination of parameters routinely used for MT preparation on clinical scanners. Indeed, Henkelman et al. go as far as deeming the use of k_for_ as „bad science‟ since it holds the pretence of being a quantitative parameter of magnetisation transfer (while not being it), opposed to the more honest, phenomenologic MTR. One needs to remain aware of the limitations and potential variability of the parameter k_for_ as defined in Eq. (7), and exercise caution in interpreting its correlations with quantitative parameters. However, mapping it offers at the very least the advantage of obtaining stunning GM-WM tissue contrast, with great potential for anatomical characterisation and segmentation (Figs. 4 and 5).

### 4.5 Correlations

A remarkable correlation between water content measured by non nuclear magnetic resonance (NMR) methods and the longitudinal relaxation time, T_1_, in biological tissue was noticed early on (Kiricuta et al., 1975; Bottomley et al., 1984; Cameron et al., 1984) and has been used by several groups to derive surrogate water content values based on T_1_ (Naruse et al., 1985; Bell et al., 1987; Bell et al., 1989; Fatouros et al., 1991; Fatouros et al., 1999). The invasive sampling of tissue, however, did not allow for an extensive characterisation of regional variation.

In this context, it is certainly interesting to note that the correlation between water content and T_1_ obtained here *in vivo* is not high at all in the WM (coefficient of below 0.3, see Fig.9a and Table 4). Indeed, myelin content has been shown to drastically contribute to the exquisite WM-GM contrast in the brain [Koenig 1991], and myelin content does not necessarily follow water content. The correlation between water content and T_1_ is better within the GM (0.6) and the extension of the range of variation to the whole brain (60-90% water content) makes it appear quite convincing (0.84). Nevertheless, as can be seen from Fig. 9a, T_1_ is not an accurate predictor of water content in an arbitrary voxel within the healthy brain. The same conclusion holds for the correlation between 1/H2O and 1/ T_1_, which would be the adequate linear correlation within a two-pool model with exchange (e.g. [Fatouros and Marmarou 1991]). Given the restricted variation in water content in the WM, for example, simply using its mean value (70%), without considering the influence of T_1_, is a valid prediction to better than 5%. However, when one is able to accurately measure the water content itself, disturbances in water homeostasis at a much lower level (1-2%) are already indicative of pathology [Shah et al. 2008]. Furthermore, spatial patterns of small water disturbances were shown to be correlated with the severity of disease in hepatic encephalopathy [Shah et al. 2008] and it is unclear for the moment whether they would be reflected by T_1_ changes. Despite all these caveats, the empirical observation remains that T_1_ and water content are generally influenced in similar ways by pathologies and, over the whole water content range in the brain, well correlated. Furthermore, T_1_ has a larger range of variation than water content, especially in the GM, and, assuming that pathological changes in T_1_ are only due to changes in water content, could in principle provide more sensitive ways to detect pathology than measurements of water content.

In fixed tissue the correlation between water content and longitudinal relaxation rate remains low in WM and becomes much lower for the GM and at whole-brain level. It is also variable between brains, probably reflecting the variability of T_1_. The same holds for 1/H2O and 1/ T_1_.

Interestingly, a higher correlation is observed between H2O and T_2_* in fixed tissue, both individually in WM and GM and for the whole brains. Whereas *in vivo* this correlation is very poor, it becomes one of the few relatively constant quantities in fixed brains (Table 4). Based on the parameters available from this study, inference of mechanisms behind the behaviour of relaxation times is sheer speculation. However, the apparently erratic behaviour of T_1_, both in value and in contrast between WM and GM, is reminiscent of effects of intermediate exchange, compared to the rather stable fast and slow exchange regimes. Exchange mediated by a membrane permeability rendered variable by several pre and post mortem factors might modulate this difference of behaviour in relaxation times between *in vivo* and post mortem tissue.

The correlation between T_1_ and T_2_ (Figs 8e and 9e), while being relatively weak *in vivo* increases in fixed tissue, and is also quite constant in the 7 brains studied here.

MTR and T_1_ appear strongly correlated *in vivo* but not so strongly correlated in fixed tissue. Since one of the main contributors to both parameters, myelin, appears little influenced by fixation, this might reflect, as mentioned before, the decrease in T_1_/ T_2_ in post mortem tissue and consequently the decrease of the contribution of direct saturation to MTR [Henkelman et al., 2001]. The good correlation found between f_bound_ and water content would be very predictable if f_bound_ were a truly quantitative parameter (the sum of free water fraction and bound proton fraction f_bound_ should give 1 if all water is MR visible and all bound protons contributed equally to MTR). Given the limited range of validity of the assumptions involved in deducing f_bound_, the high correlation found *in vivo* might also reflect the influence of direct saturation and the correlation between T_1_ and water content.

In general, a negative correlation was observed between water content and MT parameters, which can be qualitatively interpreted as meaning that less water content means higher macromolecular content. A similar observation is reported by Vavasour et al. [Vavasour et al., 2011] in MS patients as well as in healthy controls.

We mention that correlation coefficients between T_1_ and T_2_, and MTR and relaxation times are also reported in fixed tissue by Schmierer et al (Schmierer et al., 2008). The exact values are quite different from ones found in the present study, but are mostly referring to MS lesions. Also, we report here correlations obtained for whole brains; regional values might be quite different. A high value for T_1_-T_2_ correlation in fixed tissue is also reported by Schmierer et al [Schmierer et al., 2008].

## 5. Conclusions

With the increasing availability of ultra-high field systems, high-resolution imaging of the post mortem brain has gained momentum, but quantitative characterisation of fixed tissue remains scarce. We introduced here a 3D method for high-resolution mapping of water content, T_1_ and T_2_* relaxation times and semi-quantitative parameters characterising magnetisation transfer. Seven whole post mortem brains fixed by immersion in formalin were investigated with this method at 3T.

Changes in water content are highly relevant for the characterisation of disease *in vivo*, due to the high regulation of water content in healthy tissue, but they are usually in the low percentage range. Any useful method for water content measurement must cope with this challenge of very high accuracy. Very little is known about changes effected on the normal water content by fixation. We find here – to our knowledge, for the first time using MRI methods - an increased water content (73% for WM and 85% for GM) in fixed tissue compared to the *in vivo* case (69% and 81%). This is a consistent finding for all brains studied. Also, water content is found to be the quantitative parameter showing the least variability (SD of less than 3%) for the 7 brains. Whereas water regulation in fixed tissue is not anymore related to cell functionality, water is still intimately involved in osmotic equilibrium within tissue. This is perhaps the reason for the relative constancy of water content. The relaxation times T_1_ and T_2_* are shortened by fixation, with a stronger and more variable effect on T_1_. Also the contrast between WM and GM changes with fixation, with T_2_* giving often the higher contrast.

Variability might be related to pre- and post mortem history of tissue (agonal history, post mortem interval), perhaps through their influence on membrane permeability.

For practical MR imaging considerations, the high proton density contrast between WM and GM, further enhanced by T_2_* contrast, ensures that high-quality anatomical images are easily obtainable with a variety of sequences. Our mapping method is based on several multi-contrast acquisitions and offers a good basis of images for tissue segmentation.

Semi-quantitative MT parameters, such as MTR, k_for_ and f_bound_, were calculated and found to offer good to very good tissue contrast. A reduced MTR value was found in fixed tissue and assigned to some extent to a reduction of the direct saturation effect due to changes in T_1_/ T_2_ values.

Correlations between different parameters were investigated and found to be substantially different from the ones found *in vivo*. Most interestingly, the good correlation between T_1_ and water content reported many times *in vivo* is found to be quite limited for WM and diminishes substantially in fixed tissue. Instead, a good correlation between water content and T_2_* emerges, which was not found *in vivo*. In both cases, water content and bound proton fraction and T_1_ and MTR are found to be well correlated.

In conclusion, fixed brain tissue appears much more variable than living tissue, and factors other than myelin and water content appear to play a major role in the determination of relaxation times. This is not surprising given the important changes in the distribution of intra/extra cellular spaces and substances and given the large changes in membrane permeability that are induced by fixation. The results obtained on fixed brains are different from those reported on model systems [Thelwall 2006, Purea 2006] and on rat brain slices [Shepherd 2009].

Indisputably, multi-contrast imaging of fixed tissue offers huge possibilities for anatomical investigation of brain structure, which can be expected to improve even more at high fields and with advanced coil technology.

## Acknowledgements

First and foremost, the authors would like to thank the brain donors and their families. The contribution of Prof. K. Zilles and Prof. K. Amunts who provided fixed brains over the years is gratefully acknowledged. AMO expresses gratitude for help with the *in vivo* implementation of the 3D mapping method and data processing in an early stage from Mr. F. Keil, M.Sc.

